# Germ granules function as stress-responsive hubs to protect germ cell resilience

**DOI:** 10.64898/2026.10.23.761416

**Authors:** Olivia J. Gaylord, Misheel Armasanaa, Aritra Basu, Heng-Chi Lee

**Affiliations:** Department of Molecular Genetics and Cell Biology, University of Chicago, Chicago, IL, 60637, USA

## Abstract

Biomolecular condensates are dynamic, membrane-less compartments that perform critical cellular functions in the adaptation to environmental stressors. Within germ cells, germ granules are specialized condensates essential for germ cell viability, genome defense, and epigenetic inheritance. Notably, environmental stressors such as heat and starvation can disrupt genome defense and epigenetic inheritance, yet germ cell viability is often preserved. How germ cells adapt to environmental stress and recover viability remains poorly understood. Here, we investigate the impact of heat stress on the biomolecular condensates of adult germ cells in *C. elegans*. Upon heat stress, we observe extensive remodeling of the germ granule compartments, including reduced enrichment of key genome defense and epigenetic inheritance factors. Adjacent to remodeled germ granules, stress granules and P-bodies robustly assemble, suggesting these distinct condensates may functionally interact during the stress response. Indeed, we find stress granule and P-body formation within germ cells largely requires germ granule assembly factors GLH-1/VASA and MIP-1. Furthermore, we show these germ granule assembly factors and stress granule factors GTBP-1/G3BP1 and TIAR-1/TIA1, are essential for recovery of germ cell viability after heat stress. Intriguingly, stress granule factor TIAR-1/ TIA1 localizes to germ granules with and without stress, indicating germ granules are poised for the stress response. Together, our results support a model in which germ granules function as stress-responsive hubs that reorganize RNA granule networks to promote germline resilience.

## Introduction

Within the germ cells of most animals, germ granules function as biomolecular condensates that store, protect, and regulate the expression of germline and maternal mRNAs ^1–5^. In this role, germ granules are crucial for germ cell fate, genome defense, and epigenetic inheritance ^6–10^. In *C. elegans*, germ granules are multiphase condensates that comprise of multiple spatially distinct sub-compartments, including P, Z, D, and E granules together with Mutator and SIMR foci, each enriched for specialized RNA regulatory factors that support small RNA-mediated gene regulation, genome defense, and epigenetic inheritance ^11–18^. Notably, these critical RNA regulatory functions are often greatly compromised when the assembly and/or organization of germ granule sub-compartments are disrupted, and ultimately, lead to defects in gametogenesis ^11,14,16,17,19–21^. For example, loss of GLH-1/VASA helicase or LOTUS-domain protein MIP-1 leads to dramatic defects in germ granule assembly, small RNA-mediated gene regulation, and germline fertility ^19–23^. Interestingly, heat stress can also lead to changes in germ granule assembly, specifically reported for P granules and Mutator foci ^24–27^. Moreover, exposure to environmental stress also disrupts small RNA-mediated gene regulation and epigenetic inheritance ^28^. So far, to what extent environmental stress impacts germ granule sub-compartment organization and function in germ cell development remains poorly understood.

When exposed to stressful conditions, cells activate the integrated stress response pathway, leading to translation inhibition of most mRNAs and the assembly of two condensates, stress granules and P-bodies ^29–32^. Across animals and tissue types, stress granules are proposed to exert a protective role on cells, facilitating the stress response primarily by storing translationally stalled mRNAs, while P-bodies function mainly in RNA decay and regulation ^30–33^. The extent to which these two condensates coordinate RNA regulation during the stress response is an active area of investigation. In *C. elegans* adult germ cells, heat stress triggers robust assembly of stress granules marked by factors GTBP-1/G3BP1, PQN-59/UBAP2, and TIAR-1/TIA1, as well as P-bodies marked by factors CGH-1/DDX6 and CAR-1/LSM14 ^27,34–37^. Loss of stress granule factors leads to various fertility defects, including reduced brood sizes under normal growth conditions and compromised embryonic viability after heat stress, demonstrating that stress granule factors play a protective role in the *C. elegans* germline ^34,35,37^. P-bodies also perform important roles in gametogenesis, and under normal growth conditions P-body factors are observed adjacent to germ granules in *C. elegans* germ cells and embryos ^38–41^. After exposure to heat stress, both stress granules and P-bodies assemble at the adult germ cell nuclear periphery typical of germ granule localization ^34–38,42^. However, whether stress granules, P-bodies, and germ granules interact to promote germ cell survival in stressful environments remains unclear.

Here, we investigate how heat stress remodels germ granule organization in the *C. elegans* adult germline. We show that heat stress selectively remodels germ granule sub-compartments, with most sub-compartments losing their characteristic enrichment of key genome defense and epigenetic factors while the P granule sub-compartment remains largely intact. Concurrently, heat stress induces the assembly of perinuclear stress granules and enlarged P-bodies adjacent to P granules. Further, we demonstrate that robust perinuclear stress granule and P-body formation require germ granule assembly factors GLH-1 and MIP-1, revealing a previously unrecognized role for germ granules in organizing stress-responsive RNA condensates. Finally, we show that germ granule assembly factors, like stress granule factor GTBP-1 and TIAR-1, are critical for the recovery of embryonic viability following heat stress. Notably, TIAR-1 localizes to the P granule with and without stress, suggesting germ granules are poised for the stress response. Together, our findings establish germ granules as stress-responsive hubs that remodel to coordinate stress granule and P-body condensate assembly, thereby safeguarding germline resilience during stress.

## Results

### Germ granule sub-compartments are remodeled after heat stress

Environmental stress, such as heat and starvation, has been shown to disrupt small RNA-mediated gene silencing and epigenetic inheritance ^28^, functions primarily carried out by small RNA pathway factors that localize to germ granules. In *C. elegans*, germ granules are multiphase condensates, composed of sub-compartments enriched for distinct sets of small RNA factors that surround the nuclear periphery (Fig. 1A) ^11–18^. We wondered whether environmental stress leads to changes in germ granule organization. To this aim, we examined the localization of factors from each germ granule sub-compartment in adult pachytene germ cells after stress. For our study, we utilized the stress stimulus of heat stress for the induction of a rapid and robust stress response. In result, we observed a rapid reduction in granule size for most of the P, Z, D, E, and M granule sub-compartment markers after heat stress (37 °C, 15 min) compared to animals grown at normal conditions (20 °C) (Fig. 1B-F, Fig. S1A-E, and Table S1). However, not all markers from the same compartment displayed the same severity of reduction. Within the P and Z compartments, while GLH-1, PRG-1, and WAGO-4 marked granules were largely dispersed from the granule after heat stress, WAGO-1, PGL-1 and ZNFX-1 marked granules appeared similar in size to granules under normal growth conditions (Fig. 1B-C). For D and E granules, we observed a similar reduction across factors in their granule enrichment (Fig. 1D-E). Additionally, we found a reduction in Mutator foci marked by MUT-16 (Fig. 1F), supporting previously published evidence that Mutator foci are mostly dispersed from granules following heat stress ^25^. In contrast, for SIMR-1 marked SIMR foci, we observed a slight increase in puncta size after heat stress compared to no stress (Fig. 1G, Fig. S1E). These results demonstrate that sub-compartments and their specific factors can be regulated differently in response to heat stress.

**Figure 1:**
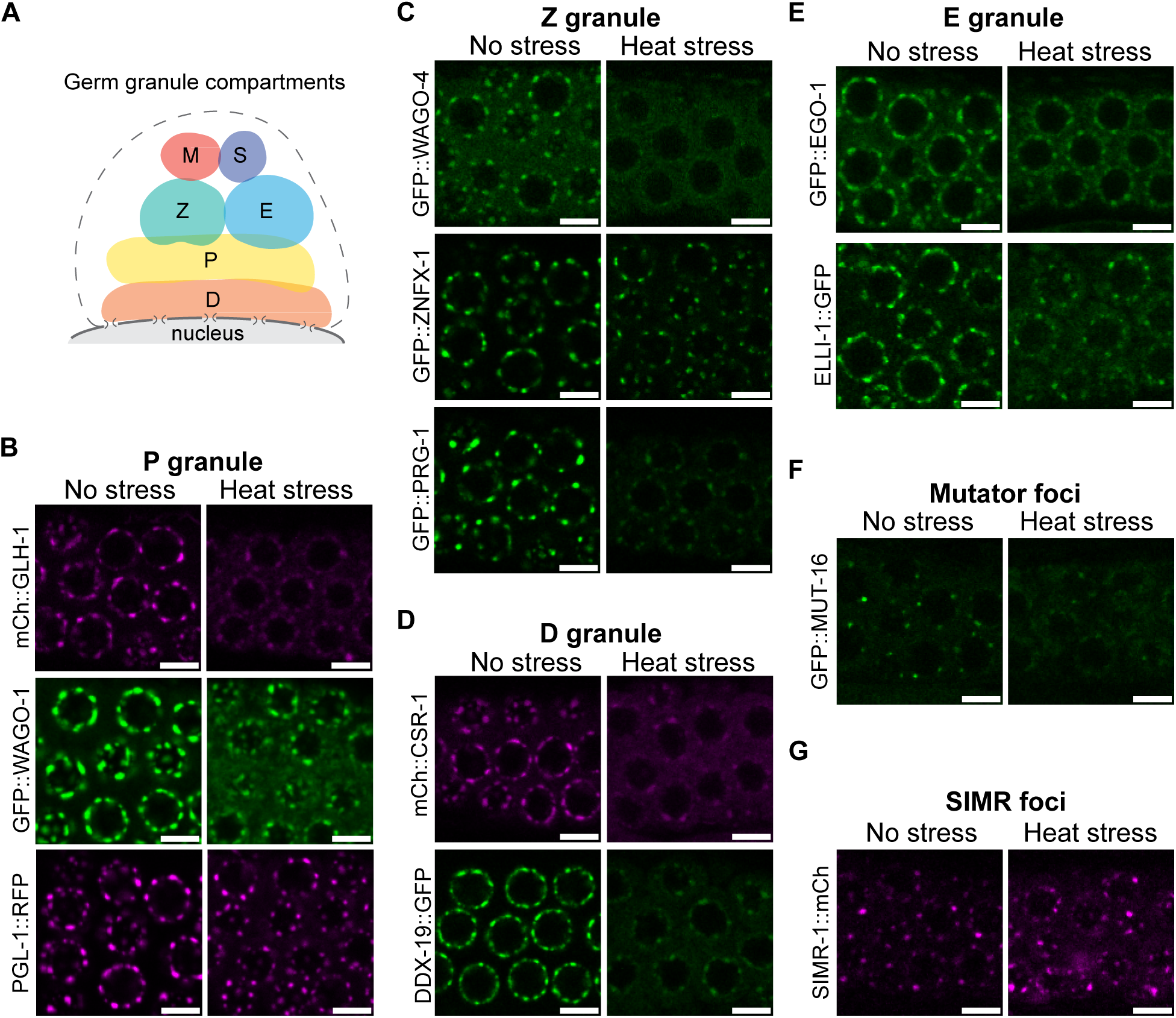
Germ granule compartments are remodeled after heat stress. (**A**) Cartoon model illustrating germ granule compartment organization in normal growth (20 °C) for day one adult pachytene germ cells. Abbreviations in cartoon P, Z, D, and E correspond to granule sub-compartments, (M) mutator foci, (S) SIMR foci. (**B** - **G**) Representative live-fluorescent images of day one adult pachytene germ cells expressing indicated granule compartment or foci markers labeled. Germlines of animals under no stress (20 °C) and heat stress (37 °C, 15m) conditions. Scale bars = 5 µm.

Germ granules are similar, but not identical in organization across the developing stages of germ cells and embryos ^43,44^. Previous work has demonstrated that PGL-1 marked P granules in oocytes rapidly decondense after heat stress ^26^. Similarly, we observed dispersal of P granules within oocytes for PGL-1, WAGO-1, and GLH-1 markers (Fig. S2A). Intriguingly, this heat stress response for PGL-1 and WAGO-1 granules in oocytes differed from pachytene germ cells where their granules remained mostly intact (Fig. 1A, Fig. S2A). While Z granule factors PRG-1 and WAGO-4 were also dispersed in oocytes under heat stress, several ZNFX-1 granules remained (Fig. S2B). For D, E, Mutator and SIMR compartment markers, we observed a reduction in oocyte granule enrichment after heat stress (Fig. S2C-F). Together, these results suggest that germ granules are rapidly remodeled following heat stress across germ cell developmental stages, with significant loss of germ granule enrichment for several compartments and small RNA factors, like MUT-16 and WAGO-4, known for their roles in genome defense and epigenetic inheritance. Nonetheless, within pachytene germ cells, several P granule factors, including PGL-1 and WAGO-1, as well as the SIMR-1 foci maintain perinuclear enrichment after acute heat stress.

### Heat stress induces perinuclear stress granules and P-bodies in germ cells

Various types of stress stimuli, including heat, oxidative, and osmotic stresses, have been shown to trigger formation of stress granules and P-bodies in the germline ^24,34,35,37,45^. Notably, these two stress-induced condensates have been shown to form in germ cells near germ granules ^24,34,35,37,45^. In follow up of these reports, we examined the localization of conserved stress granule components GTBP-1/G3BP1, PQN-59/UBAP2, and TIAR-1/TIA1 in the young adult germline after heat stress. Under normal growth conditions, GTBP-1 and PQN-59 localized evenly throughout the germline cytoplasm (Fig. 2A). Distinct from GTBP-1 and PQN-59, TIAR-1 showed nuclear and cytoplasmic localization under normal growth, as well as some granular enrichment at the nuclear periphery (Fig. 2A). When animals were exposed to heat stress, similar to previous reports, we observed GTBP-1, PQN-59, and TIAR-1 formed robust granules surrounding the pachytene germ cell nuclei (Fig. 2A) ^34,35,37,42^. In addition to these stress granules, TIAR-1 also maintained nuclear localization (Fig. 2A). These localization differences between stress granule factors may reflect their distinct roles during germ cell development and the stress response ^35,37,42^.

**Figure 2:**
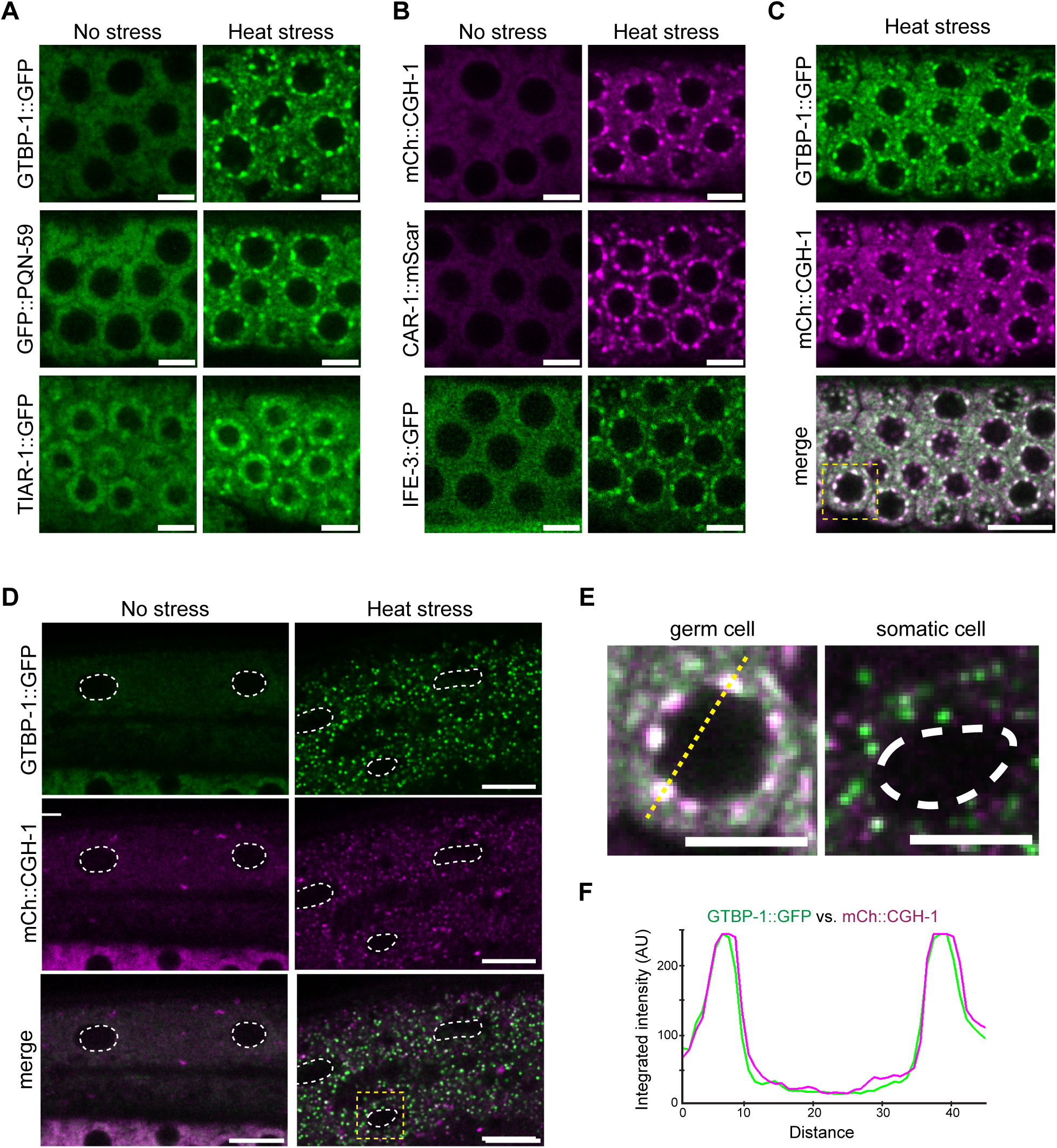
Distinct patterns of heat stress-induced stress granule and P-body assembly between the germline and soma. (**A** and **B**) Representative live-fluorescent images of day one adult pachytene germ cells expressing indicated stress granule markers for no stress (20 °C) or heat stress (37 °C, 15 min) conditions. Scale bars = 5 µm. (**C** and **D**) Representative live-fluorescent images of animals co-expressing indicated markers under heat stress (37 °C, 15m) in adult pachytene germ cells (C) and no stress (20 °C) and heat stress (37 °C, 30m) in somatic cells (D). White dashed lines mark individual somatic cell nuclei. Scale bars = 10 µm. (**E**) Yellow dashed boxes in A and B outline the merged image shown enlarged. Scale bars = 5 µm. (**F**) Line scans display the integrated fluorescent intensity corresponding to the yellow dashed line across the germ cell nucleus shown in (E) for the indicated markers labeled in green and magenta.

Stress granules share several components with P-bodies, and exposure to stress can also enhance P-body assembly ^30,33,46^. Additionally, under normal growth conditions, P-bodies have been found to interact with germ granules in the adult *C. elegans* germline ^39^. Therefore, we next examined the localization of conserved P-body components CGH-1/DDX6, CAR-1/LSM14, and IFE-3/EIF4E after exposure to heat stress. In the young adult germ cells of animals under normal growth conditions, CGH-1, CAR-1, and IFE-3 localized evenly throughout the cytoplasm (Fig. 2B). At this early adult stage, P-bodies are not yet robustly assembled at the site of germ granules. Nonetheless, when animals were exposed to heat stress, CGH-1, CAR-1, and IFE-3 assembled into granules surrounding the perinucleus of germ cells (Fig. 2B).

Based on the enrichment of both stress granules and P-bodies surrounding the nuclear periphery of germ cells under heat stress, we examined whether stress granules and P-bodies form at overlapping perinuclear sites. In pachytene germ cells under heat stress, GTBP-1 marked stress granules and CGH-1 marked P-bodies assembled at highly overlapping sites surrounding the germ cell nuclei (Fig. 2C). Closer inspection of the relationship between stress granules and P-bodies surrounding an individual pachytene nucleus in heat stressed animals showed near perfect overlap of stress granules with P-bodies (Fig. 2D-E). Notably, this formation of both stress granules and P-bodies closely surrounding the nuclear periphery was not observed in somatic cells under heat stress (Fig. 2E). Rather, in response to heat stress, stress granules marked by GTBP-1 formed evenly throughout the cytoplasm of somatic cells (Fig. 2F). Additionally, CGH-1 marked P-bodies were not as robustly induced in somatic cells compared to germ cells after heat stress (Fig. 2F), likely due to CGH-1 primary expression in the germline and early embryos ^38^. Together, our results suggest stress granules and P-bodies, in response to heat stress, form highly overlapping granules at the nuclear periphery of germ cells. Moreover, this localization pattern, appears to mainly occur within germ cells and resembled the position of germ granules ^1,18^.

### Stress granules and P-bodies form at the cytoplasmic surface of germ granules

Following heat stress, we observed the induction of stress granule and P-body formation at the germ cell perinucleus where germ granules are typically found. We wondered whether these three types of condensates associated together or formed distinct granules surrounding the nuclear periphery within the adult germline. To this aim, we examined animals co-expressing markers for P granules, the germ granule sub-compartment we found remained largely intact after heat stress, with stress granules and/or P-bodies under normal or heat stress conditions.

First, we compared the localization of stress granule factors GTBP-1 and PQN-59, as well as P-body factor CAR-1, with P granules in adult pachytene germ cells (Fig. 3A-C). As expected under normal growth, these stress granule and P-body factors localized evenly throughout the cytoplasm (Fig. 3A-C). After exposure to heat stress, we observed the robust assembly of GTBP-1, PQN-59, and CAR-1 granules adjacent to the P granules (Fig. 3A-C). Upon closer inspection, we found GTBP-1 and PQN-59 marked stress granules, as well as CAR-1 marked P-bodies, formed partially overlapped with but on the cytoplasmic surface of P granules (Fig. 3A-F). More specifically, the integrated intensity of P granules showed peaks slightly inside of those for both stress granule and P-body markers (Fig. 3D-F). Together, these results demonstrate heat stress-induced stress granule and P-bodies assemble at similar sites as P granules, occupying spatially distinct but partially overlapped compartments.

**Figure 3:**
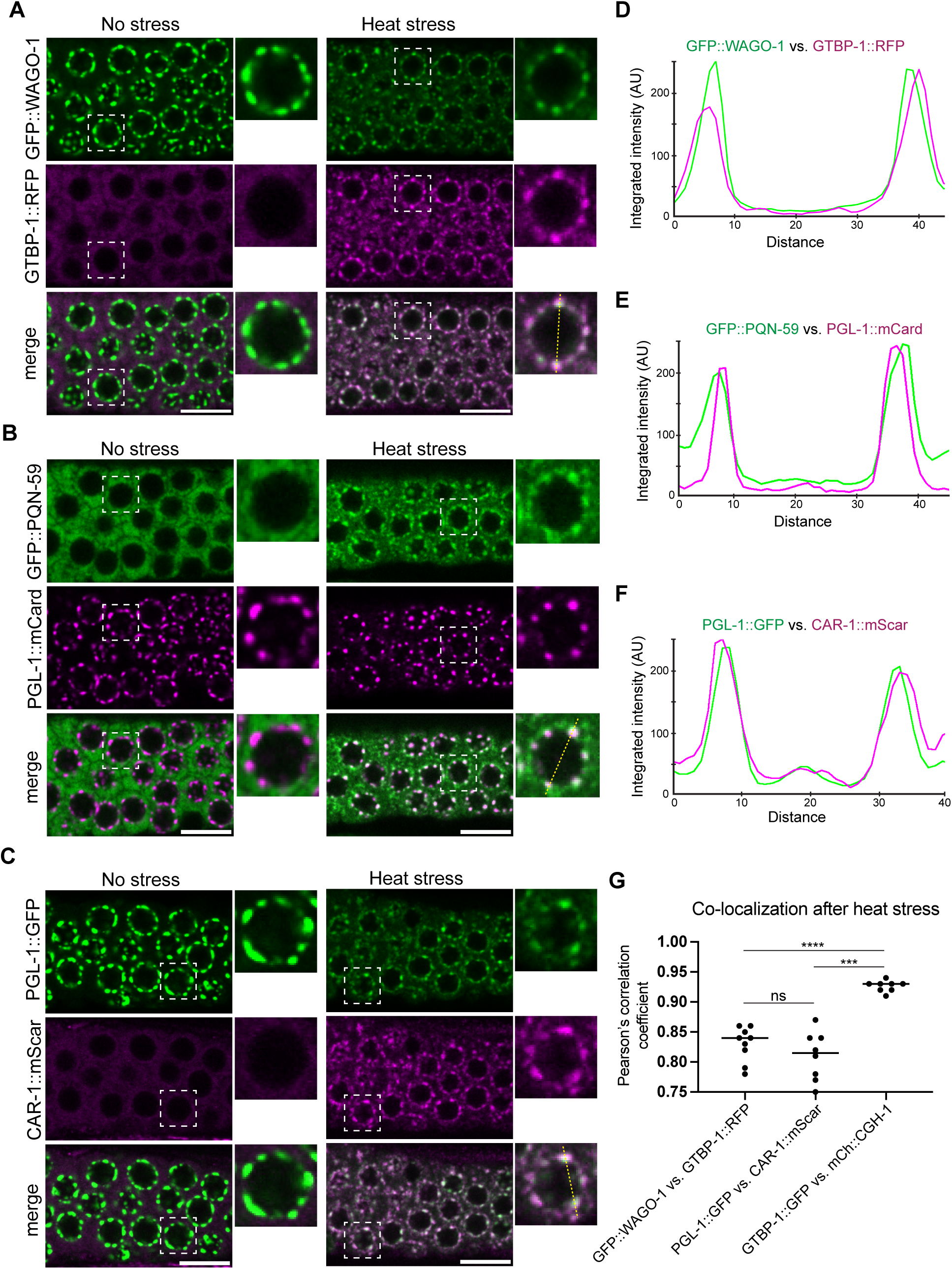
Stress granules and P-bodies assemble at P granules after heat stress. (**A** - **C**) Representative live-fluorescent images of pachytene germ cells co-expressing the indicated markers in no stress (20 °C) or heat stress (37 °C, 15 min) conditions. Dashed white boxes outline the individual germ cell nucleus shown in the enlarged image. Scale bars = 10 µm. (**D** - **F**) Line scans display the integrated fluorescent intensity corresponding to the yellow dashed line across the nucleus shown in (A to C) for the indicated markers labeled in green and magenta in heat stress. (**G**) Quantification of co-localization between the indicated fluorescent proteins for animals exposed to heat stress (37 °C, 15 min). Each data point represents the Pearson’s R-value showing the degree of co-localization between two fluorescence channels. The solid black line indicates the median value. Each data point represents independent gonad images. *N =* 8-9 gonads. Two-tailed p value was calculated using Mann-Whitney-Wilcoxon test. ***, p<0.001; ****, p<0.0001; ns, no significance.

To further explore whether perinuclear stress granules and P-bodies occurred mainly at sites with P granules, we compared the localization of stress granule and P-bodies to P granules in oocytes under heat stress. In oocytes, WAGO-1 and PGL-1 marked P granules were greatly reduced after heat stress as previously reported for P granules ^26,27^, while stress granules formed throughout the cytoplasm without robust perinuclear enrichment (Fig. S3A-B). In very few early oocytes under heat stress, we observed several perinuclear WAGO-1 marked P granules that coincided with GTBP-1 marked stress granules (Fig. S3C). Similarly in early oocytes under heat stress, we observed several perinuclear PGL-1 marked P granules that coincided with CAR-1 marked P-bodies (Fig. S4A-B). These results suggest the formation of heat-induced perinuclear stress granules and P-bodies within germ cells primarily occur at sites with perinuclear P granules.

Altogether, our observations suggested stress granules and P-bodies highly overlap at the cytoplasmic side of P granules in germ cells after heat stress. To support this, we performed co-localization analysis to quantify the pair-wise correlation of P granules, stress granules, and P-bodies under heat stress. As expected from our observations, we found the correlation values of WAGO-1 marked P granules to GTBP-1 marked stress granules were comparable to those of PGL-1 marked P granules to CAR-1 marked P-bodies (Fig. 3G). Also consistent with our observations, the correlation values of GTBP-1 marked stress granules to CGH-1 marked P-bodies were significantly higher compared to either stress granule or P-body marker co-localization with P granule markers (Fig. 3G). This suggests stress granules, P-bodies, and P granules form closely associated but separate compartments under heat stress.

Distinct from GTBP-1 and PQN-59, stress granule factor TIAR-1 has been noted in previous reports to localize in the nucleus and perinucleus of germ cells under normal growth ^35,42^. In addition to cytoplasmic and nuclear localization under normal growth, we observed TIAR-1 enrichment within granules that largely overlapped with PGL-1 marked P granules (Fig. 4A-B). After exposure to heat stress, TIAR-1 was further enriched at sites overlapping with perinuclear P granules (Fig. 4A and C). Furthermore, we found TIAR-1 marked granules were often adjacent, and in some cases, highly overlapped with ZNFX-1 marked Z granules, comparable to the known relationship of the P and Z compartments within germ granules (Fig. S5A-C). This supported TIAR-1 localization to the P granule sub-compartment. Additionally, we compared the localization of TIAR-1 to stress granule and P-body markers, GTBP-1 and CGH-1, respectively. We found both GTBP-1 and CGH-1 granules assembled highly overlapped with TIAR-1 granules, but not all TIAR-1 granules had corresponding GTBP-1 or CGH-1 granules (Fig. 4B-C). Additionally, for several granules, TIAR-1 integrated intensity peaks were slightly inside those for GTBP-1 and CGH-1 markers, similar to our observations of these markers relative to P granules (Fig. 4D-G).

**Figure 4:**
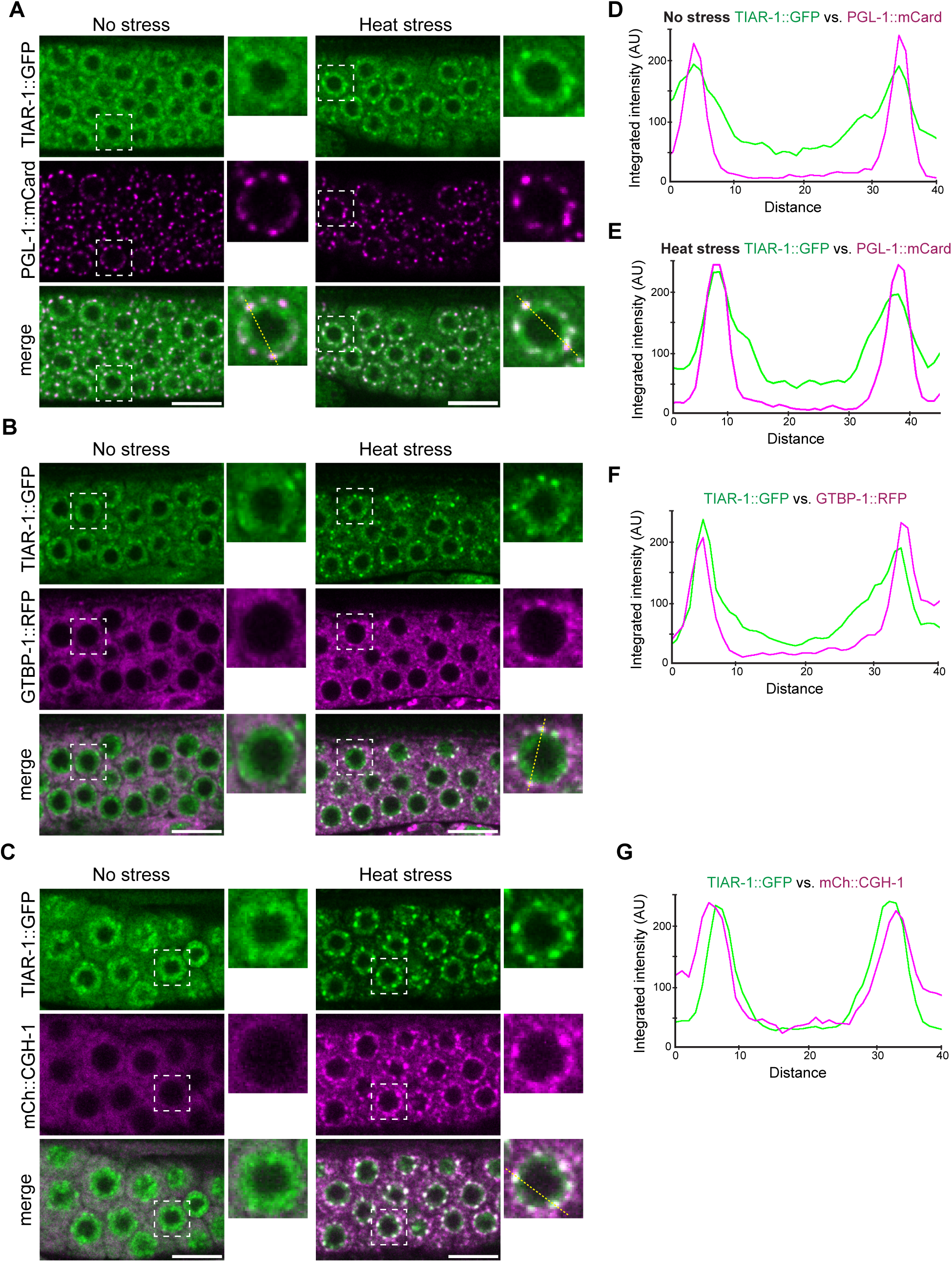
Stress granule factor TIAR-1 localizes to P granules under normal growth and heat stress. (**A** - **C**) Representative live-fluorescent images of pachytene germ cells co-expressing the indicated markers in no stress (20 °C) or heat stress (37 °C, 15 min) conditions. Dashed white boxes outline the individual germ cell nucleus shown in the enlarged image. Scale bars = 10 µm. (**D** - **G**) Line scans display the integrated fluorescent intensity corresponding to the yellow dashed line across the nucleus shown in (A to C) for the indicated markers labeled in green and magenta in no stress (D) or heat stress (E to G) conditions.

Together, our findings demonstrate that following heat stress in adult germ cells, stress granules and P-bodies closely assembly at the cytoplasmic surface of P granules. Unlike stress granule factors GTBP-1 and PQN-59, our observations position stress granule factor TIAR-1 within P granules in both normal growth and heat stress, likely indicative of the distinct and varied roles of these factors within germ cells.

### Stress granule and P-body assembly is largely independent in germ cells after heat stress

Based on the high co-localization and near perfect overlap of stress granules and P-bodies at the nuclear periphery of germ cells after heat stress, we next asked whether stress granule and P-body assembly occur independently of each other. While these granules are widely proposed to form independent but highly interactive condensates after stress ^46–48^, several studies suggest stress granule assembly may also require P-body assembly factors ^49,50^. Toward the latter case, P-bodies have been shown to promote stress granule assembly in yeast ^49^. Additionally, in mammalian cells, P-body factor DDX6/CGH-1 was found to promote the assembly of stress granules during oxidative stress ^50^. Together, these studies highlight a complex cell- and stress-context dependent and potential interactive assembly of P-bodies and stress granules.

To test whether stress granules assemble independently of P-bodies in the *C. elegans* germline after heat stress, we performed RNAi knockdown of the core P-body assembly factors CGH-1 and CAR-1 in animals co-expressing stress granule marker GTBP-1 and P-body marker CGH-1. In *cgh-1* knockdown, we observed loss of mCh::CGH-1 signal, indicating successful knockdown (Fig. 5A-B, Fig. S6A-B, Fig. S7A-B). In *car-1* knockdowns, we observed reduced P-body formation, marked by CGH-1, after heat stress (Fig. 5A-B, Fig. S6A-B, Fig. S7A-B). This observation is consistent with the critical role of CAR-1 in P body assembly ^39^. However, under heat stress, GTBP-1 marked stress granules robustly assembled at the nuclear periphery of germ cells in empty vector, *cgh-1,* and *car-1* knockdown animals (Fig. 5A-B, Fig. S6A-B, Fig.S7A-B). These findings suggest stress granule assembly in the pachytene germ cells after heat stress is largely independent of P-body assembly.

**Figure 5:**
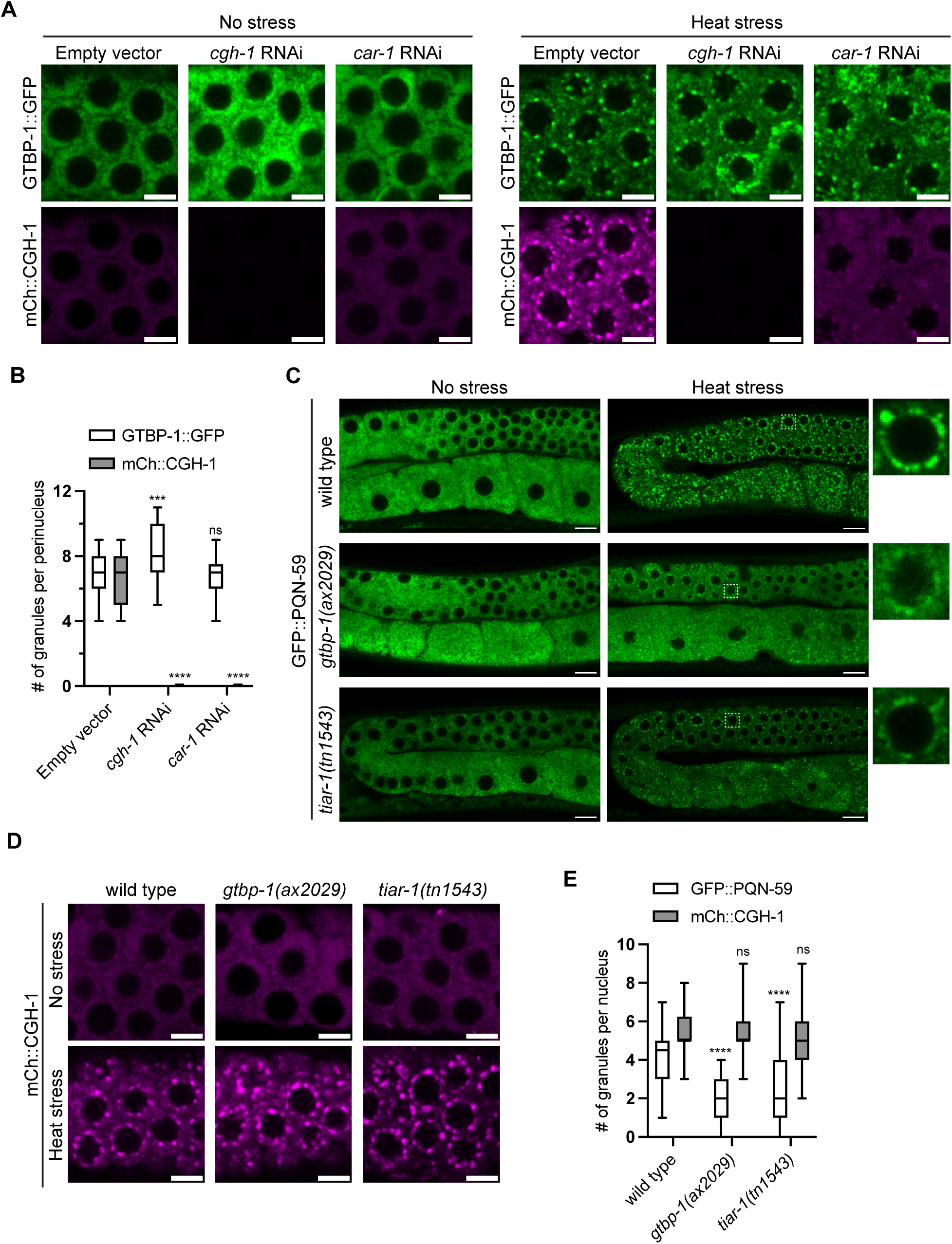
Largely independent assembly of perinuclear stress granules and P-bodies in germ cells after heat stress. (**A**) Representative live-fluorescent images of day one adult pachytene germ cells co-expressing the indicated markers for control empty vector, *cgh-*1, or *car-1* RNAi conditions for no stress (20 °C) (left) and heat stress (37 °C, 15m) (right) conditions. Scale bars = 5 µm. (**B** and **E**) Boxplots showing the number of granules per germ cell perinucleus for indicated markers for animals exposed to heat stress under the labeled RNAi knockdown (**B**) or mutant (**E**) conditions. Quantified for 5-6 cells from N=8-12 independent germlines. (**C**) Representative live-fluorescent images of day one adult germline expressing GFP::PQN-59 in wild type or indicated mutants for no stress (20 °C) (left) and heat stress (37 °C, 15m) (right). White dashed boxes outline pachytene germ cell shown enlarged. Scale bars = 10 µm. (**D**) Representative live-fluorescent images of day one adult pachytene germ cells expressing P-body marker mCh::CGH-1 in wild type or indicated mutant in no stress (20 °C) (left) and heat stress (37 °C, 15m) (right) conditions. Scale bars = 5 µm. For all boxplots, the line indicates the median value, the box indicates the first and third quartiles, and the whiskers indicate the min and max values. Two-tailed p values were calculated using Mann-Whitney-Wilcoxon test. ***, p<0,001; ****, p<0.0001; ns, no significance.

We next examined whether P-bodies form independently of stress granule assembly factors after heat stress. To this aim, we evaluated heat stress-induced P-body formation in *gtbp-1(ax2909)* and *tiar-1(tn1543)* null mutants. First, we found stress granule formation, marked by PQN-59, was largely compromised in both *gtbp-1(ax2909)* and *tiar-1(tn1543)* mutants compared to wild type after heat stress, consistent with roles for GTBP-1 and TIAR-1 in promoting stress granule assembly (Fig. 4C). Specifically, while the size of PQN-59 marked stress granules were similar, both the number per germ cell and mean intensity were significantly reduced in *gtbp-1(ax2909)* and *tiar-1(tn1543)* mutants compared to wild type after heat stress (Fig. 4C and 4E, Fig. S7C-D). In these mutants with compromised stress granule assembly, we observed similar heat-stress induced P-body formation, marked by CGH-1, as wild type for pachytene germ cells (Fig. 4D). Both the number per perinucleus and mean intensity of CGH-1 marked P-bodies were similar between wild type, *gtbp-1(ax2909)* and *tiar-1(tn1543)* mutants after heat stress, while the average size of CGH-1 marked P-bodies was slightly increased in *gtbp-1(ax2909)* and *tiar-1(tn1543)* mutants (Fig. 4E and Fig. S7E-F). Consistent with these results, we similarly observed robust heat stress-induced mCh::CGH-1 P-body formation in germ cells following knockdown of endogenously tagged GTBP-1::GFP and TIAR-1::GFP by *gfp* RNAi in co-marked backgrounds (Fig. S6A-D). Additionally, we found CAR-1 marked P-bodies assembled in germ cells after heat stress in *tiar-1(tn1543)* mutants comparable to wild type (Fig. S8A-B). Taken together, these results suggest P-bodies maintain perinuclear assembly in stress granule compromised backgrounds in response to heat stress.

Since stress granule assembly, as marked by PQN-59, was not completely blocked in either *gtbp-1(ax2909)* or *tiar-1(tn1543)* mutants after heat stress, we also examined the relationship between these factors in their granule assembly. After heat stress, we found neither GTBP-1 nor TIAR-1 were required for the assembly of the other at the germ cell perinucleus, suggesting single *gtbp-1(ax2909)* and *tiar-1(tn1543)* mutant animals are only partially compromised in stress granule formation (Fig. S8C-F). This suggests these core stress granule assembly factors can still form partial granules in the absence of the other during acute heat stress.

Overall, although stress granules and P-bodies highly co-localize after acute heat stress around the perinucleus of pachytene germ cells, we find the assembly of these RNA granules is largely independent under these conditions.

### Germ granules promote perinuclear stress granule and P-body assembly

Based on our results that stress granules and P-bodies assemble largely independently and adjacent to perinuclear P granules in germ cells after heat stress, we hypothesized that germ granules may serve as a docking site for their assembly. If so, disrupting P granule assembly should compromise stress granule and P-body formation at the nuclear periphery after heat stress. To test this, we examined stress granule and P-body formation after heat stress in the *glh-1(uoc1)* or *mip-1(uae1)* null mutants, two well characterized germ granule assembly factors important for P and Z compartments ^19–21,23,51,52^. More specifically, we evaluated stress granule formation in animals co-expressing PQN-59 and P granule factor PGL-1 markers, and P-body formation in animals co-expressing CGH-1 and Z granule factor WAGO-4 markers. As expected without stress, we observed dramatic reduction in the overall number and size of both PGL-1 marked P granules and WAGO-4 marked Z granules in *glh-1(uoc1)* and *mip-1(uae1)* mutants compared to wild type (Fig. 6A-D and Fig. S9A-B). While the mean intensity of PGL-1 marked P and WAGO-4 marked Z granules was significantly decreased in *glh-1(uoc1)* mutants, only WAGO-4 marked Z granules showed a decrease in *mip-1(uae1)* mutants as these mutants maintain several bright enlarged PGL-1 marked P granules (Fig. S9C-D). These results are consistent with reported function of GLH-1 and MIP-1 and reflect their slightly distinct roles in germ granule assembly ^19–21,53^.

**Figure 6:**
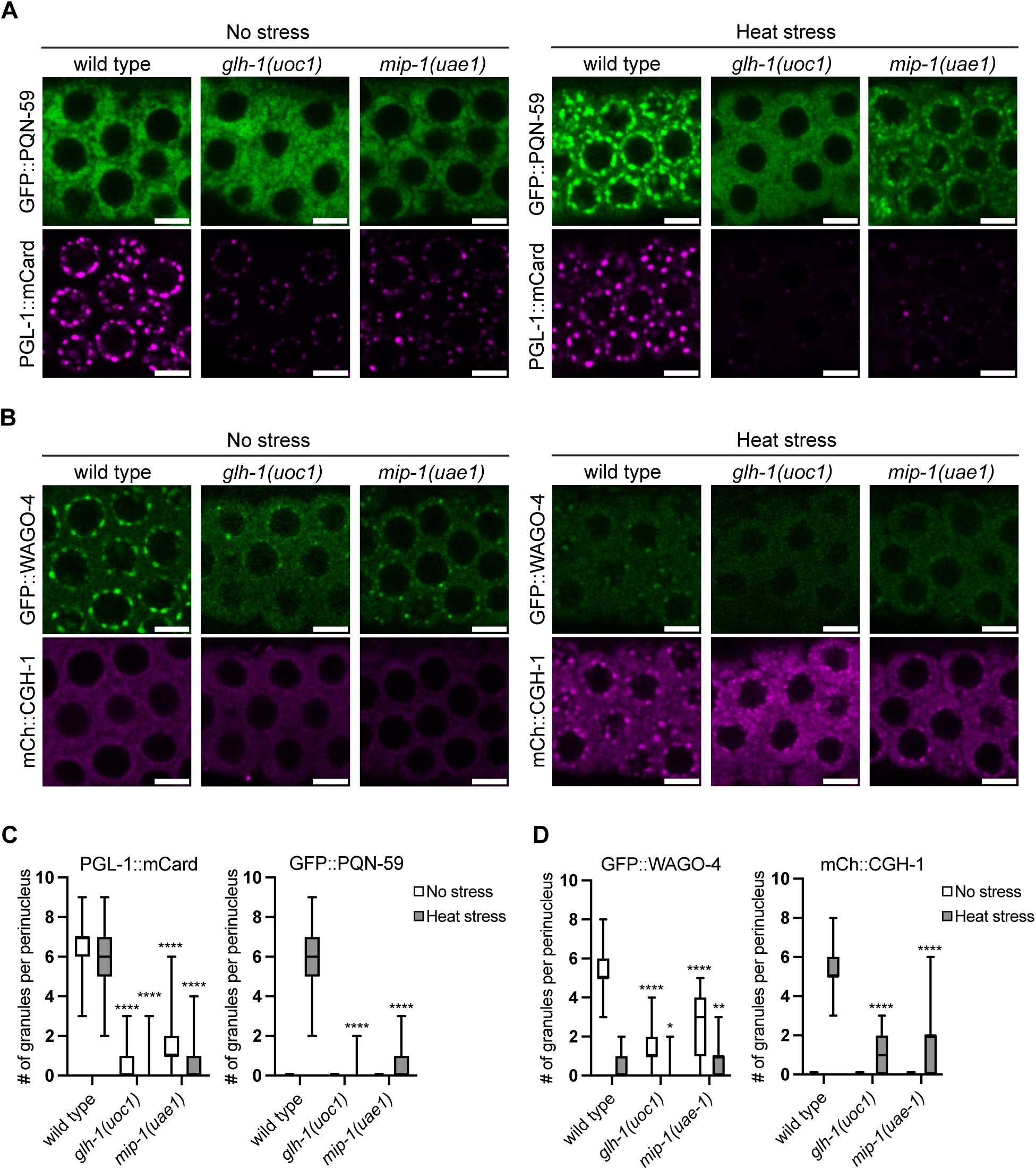
Germ granule assembly factors promote perinuclear stress granule and P-body formation after heat stress. (**A** and **B**) Representative live-fluorescent images of day one adult pachytene germ cells co-expressing GFP::PQN-59 and PGL-1::mCard (A) or GFP::WAGO-4 and mCh::CGH-1 (B) for wild type, *glh-1(uoc1)*, and *mip-1(uae1)* mutants in no stress (20 °C) (left) and heat stress (37 °C, 15m) (right) conditions. Scale bars = 5 µm. (**C** and **D**) Boxplots showing quantification of the number of granules per germ cell perinucleus for experiments shown in (A and B). No stress (white boxes), Heat stress (gray boxes). Quantified for 5-6 cells from N=8-15 independent germlines. For all boxplots, the line indicates the median value, the box indicates the first and third quartiles, and the whiskers indicate the min and max values. Two-tailed p values were calculated using Mann-Whitney-Wilcoxon test. *, p<0.05; **, p<0.01; ****, p<0.0001.

Following exposure to heat stress, as expected for wild type animals we observed robust formation of PQN-59 marked stress granules adjacent to P granules, as well as CGH-1 marked P-bodies (Fig. 6A-B). In contrast, *glh-1(uoc1)* and *mip-1(uae1)* mutants showed dramatically reduced formation of both PQN-59 marked stress granules and CGH-1 marked P-bodies (Fig. 6A-B). Consistent with these observations, the number, size, and intensity of PQN-59 granules were significantly reduced in *glh-1(uoc1)* and *mip-1(uae1)* mutants compared to wild type (Fig. 6A and Fig. S9E-F). In addition, the number and size of CGH-1 marked P-bodies were reduced in *glh-1(uoc1)* and *mip-1(uae1)* mutants compared to wild type, while their mean intensity was similar (Fig. 6A and Fig. S9G-H). Concurrently, we observed a near absence of PGL-1 and WAGO-4 marked P and Z granules in *glh-1(uoc1)* and *mip-1(uae1)* mutants compared to their number and size in wild type after heat stress (Fig. 6A-D and Fig. S9A-B). Together, these results demonstrate that germ granule assembly factors facilitate heat-stress induced perinuclear stress granule and P-body assembly.

To further examine the roles of P granules in acute heat stress-induced stress granule and P-body formation, we evaluated additional markers in either mutant or *glh-1* and *mip-1* RNAi knockdown animals. First, we examined stress granule factor TIAR-1 and P-body factor CAR-1 markers in germ granule assembly factor *mip-1(uae1)* mutants. Under non-stress and heat stress, we observed reduced TIAR-1 marked stress granules concurrent with reduced PGL-1 marked P granules in *mip-1(uae1)* mutants compared to wild type (Fig. S10A-B). For CAR-1 marked P-bodies, we also found reduced assembly in *mip-1(uae1)* mutants compared to wild type (Fig. S10C-D). Additionally, we performed *mip-1* and *glh-1* RNAi knockdowns in animals co-expressing a combination of stress granule, P granule, or P-body markers. First, in animals co-expressing WAGO-1 marked P granules and GTBP-1 marked stress granules, we observed P granule defects consistent with successful knockdown of *glh-1* and *mip-1* (Fig. S11A-B). Under heat stress, we observed reduced formation of GTBP-1 stress granules in both *glh-1* and *mip-1* knockdowns compared to empty vector animals (Fig. S11A-B). Similarly, in animals co-expressing GTBP-1 marked stress granules and CGH-1 marked P-bodies, we observed reduced stress granule and P-body assembly in both *glh-1* and *mip-1* knockdowns compared to empty vector animals under heat stress (Fig. S11C-D).

Overall, we find P granule assembly factors, including GLH-1 and MIP-1, promote stress granule and P-body assembly at the perinucleus of germ cells following heat stress. Our results suggest that P granules may serve as docking sites or seeds for both stress granule and P-body assembly in germ cells.

### Germ granule assembly factors protect germ cell viability after heat stress

The stress response, including stress granule and enhanced P-body formation, is critical for cellular and organismal recovery from environmental stressors ^35,45^. Since we found heat-induced stress granule and P-body formation is largely compromised in germ granule assembly mutants, we wondered if germ granule assembly factors contribute to the recovery of germ cell viability after heat stress. To test this, we utilized a heat stress germ cell viability assay established in a previous report ^35^. In this assay, young adult animals are exposed to a longer, lower intensity heat stress (31 °C, 3 hours), then moved back to normal growth temperature (20 °C) where embryonic lethality is measured for embryos laid over three recovery time windows ^35^. Since the adult hermaphrodite germline functions as a continuous assembly line of female gamete production, embryos laid in later time windows are produced from germ cells that were less mature during the heat stress. The published work found that stress granule factor *tiar-1* mutant animals suffer a higher percentage of embryonic lethality and take a longer time to recover embryo viability compared to wild type animals ^35^, suggesting the stress resilience and recovery of germ cell viability depends on stress granule factors.

To further investigate the role of stress granule factors in germline resilience to heat stress, we examined the recovery of germ cell viability in *tiar-1(tn1543)* and *gtbp-1(ax2029)* mutant animals which both showed a strong reduction in PQN-59 marked stress granules compared to wildtype after heat stress (31 °C, 3h) (Fig. 7A). Compared to wild type, both *tiar-1(tn1543)* and *gtbp-1(ax2029)* mutant animals showed increased embryonic lethality for embryos laid in the first recovery group after 31 °C 3h heat stress (group I in Fig. 7B). For embryos laid at longer recovery times after heat stress, both *tiar-1(tn1543)* and *gtbp-1(ax2029)* mutants also displayed delayed recovery of embryo viability compared to wild type (group II and III in Fig. 7B). Notably, *tiar-1(tn1543)* mutant animals displayed higher embryonic lethality in all groups compared to *gtbp-1(ax2029)* mutants, suggesting TIAR-1 may perform a more critical role in germ cell recovery (Fig. 7B). Additionally, under non-stress conditions, *tiar-1(tn1543)*, but not *gtbp-1(ax2029)*, animals exhibited an increase in embryonic lethality, potentially due to additional roles of TIAR-1 in germline maintenance or embryonic development (Fig. 7B). Our results for *tiar-1(tn1543)* and *gtbp-1(ax2029)* mutant animals support those previously published for *tiar-1* mutants ^35^, reinforcing that stress response factors are critical for germ cell recovery and embryo viability after heat stress.

**Figure 7:**
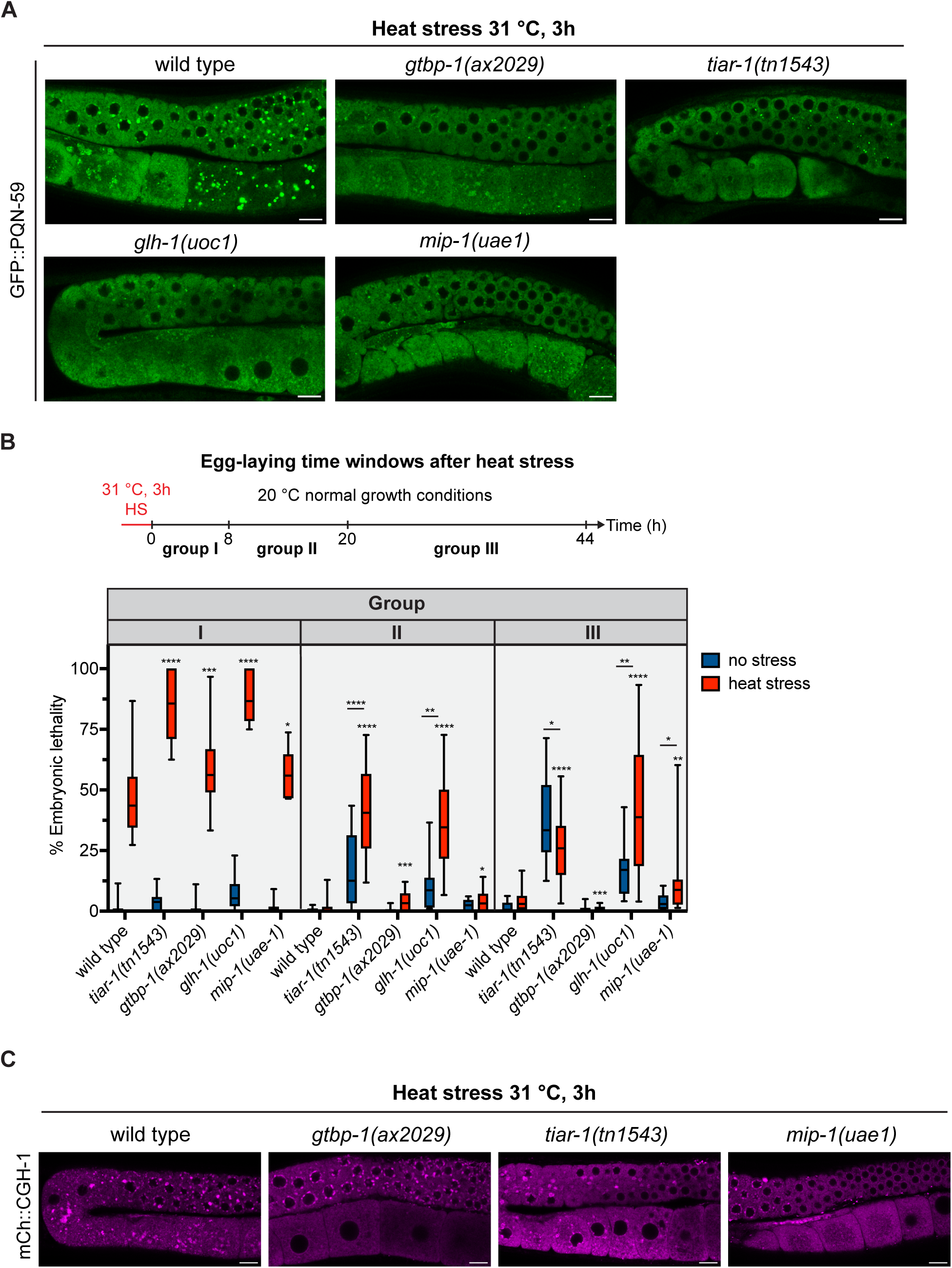
Germ granule assembly factors promote recovery of germ cell viability after heat stress. (**A** and **C**) Representative live-fluorescent images of day one adult germlines expressing indicated markers in indicated background exposed to heat stress (31 °C, 3h). Scale bars = 10µm. (**B**) Top, experiment timeline outlining heat stress (31 °C, 3 h HS) and transfer of animals to recover at normal growth conditions 20 °C. During recovery at 20 °C, animals were single picked and transferred to new plates at indicated timepoints in hours (h). Living hatched worms and unhatched embryos were quantified 24h after transferring the parent to new plate. Groups indicate the timing of embryos laid after heat stress. Bottom, boxplots showing the percentage of embryonic lethality quantified as the number of unhatched or dead embryos divided by the total sum of unhatched embryos and hatched progeny. Strains and conditions indicated. N=10-20 day one adult animals. For the boxplots, the line indicates the median value, the box indicates the first and third quartiles, and the whiskers indicate the min and max values. Two-tailed p values were calculated using Mann-Whitney-Wilcoxon test. *, p<0.05; **, p<0.01; ***, p<0.001; ****, p<0.0001; ns, no significance.

Next, we tested the role of germ granule assembly factors GLH-1 and MIP-1 in germline resilience to heat stress. Like both stress granule factor mutants, we found reduced PQN-59 marked stress granules in *mip-1(uae1)* and *glh-1(uoc1)* mutant germlines compared to wild type after heat stress (31 °C, 3h), supporting the role of germ granules in promoting stress granule assembly (Fig. 7A). Under non-stress conditions, *glh-1(uoc1)*, but not *mip-1(uae1)*, animals exhibited a modest increase in embryonic lethality, which may reflect the critical roles of GLH-1 in germline maintenance and fertility (Fig. 7B). Like the stress granule factor mutants, we found that both *glh-1(uoc1)* and *mip-1(uae1)* mutant animals showed increased embryonic lethality compared to wild type for embryos laid in the first group following heat stress (group I in Fig. 7B). Additionally, both *glh-1(uoc1)* and *mip-1(uae1)* mutant animals displayed a further delayed recovery of embryo viability after heat stress when compared to wild type (group II and III in Fig. 7B). Together, our findings suggest germ granule assembly factors GLH-1 and MIP-1, like conserved stress response factors TIAR-1 and GTBP-1, promote germ cell resilience and recovery of viability after heat stress.

Notably, unlike *gtbp-1(ax2029)* mutant animals, *tiar-1(tn1543), glh-1(uoc1)*, and *mip-1(uae1)* mutant animals continued to show higher levels of embryonic lethality compared to wild type (group III in Fig. 7B). We wondered whether these mutants showed further defects in P-body formation. Following 31 °C 3h heat stress, while *gtbp-1(ax2029)* mutant germlines showed CGH-1 marked P-body formation comparable to wild type, *tiar-1(tn1543)* and *mip-1(uae1)* mutant germlines showed reduced P-body formation (Fig. 7C). Our results for reduced CGH-1 granule formation after heat stress in *tiar-1(tn1543)* mutant germlines support those observed in a previous report ^35^. Taken together this suggests germ granules, including assembly factors and P granule localized TIAR-1, promote both stress granule and P-body formation during this longer, lower intensity heat stress.

Following from this result, we next examined whether germ granules serve as the docking sites or seeds for stress granules and P-bodies in the germline under this heat stress regimen. Interestingly, while most germ granule sub-compartment markers were mostly reduced following heat stress (31 °C, 3h), P granule markers GLH-1, WAGO-1, and PGL-1 all showed fewer but enlarged granules, supporting P granules as possible docking sites (Fig. S12A-F and Table S1). To test this, we examined stress granule and P-body formation relative to P granules in wild type and *mip-1(uae1)* mutant animals. First, in animals co-expressing stress granule factor TIAR-1 and P granule marker PGL-1, we observed enlarged and highly overlapped TIAR-1 and PGL-1 granules throughout the germline under heat stress compared to normal growth (Fig. 8A). In contrast, *mip-1(uae1)* mutant germlines displayed strongly reduced P granule and TIAR-1 stress granule assembly compared to wildtype under heat stress (Fig. 8A). Moreover, in animals co-expressing P-body factor CAR-1 and P granule marker PGL-1, we observed enlarged and highly overlapped CGH-1 and PGL-1 granules throughout the germline under heat stress compared to non-stress (Fig. 8B). In contrast, *mip-1(uae1)* mutant germlines displayed strongly reduced P granule and CAR-1 P-body assembly compared to wildtype under heat stress (Fig. 8B). Together, this supported our hypothesis that P granules may serve as seeds for stress granule and P-body assembly in germlines under heat stress.

**Figure 8:**
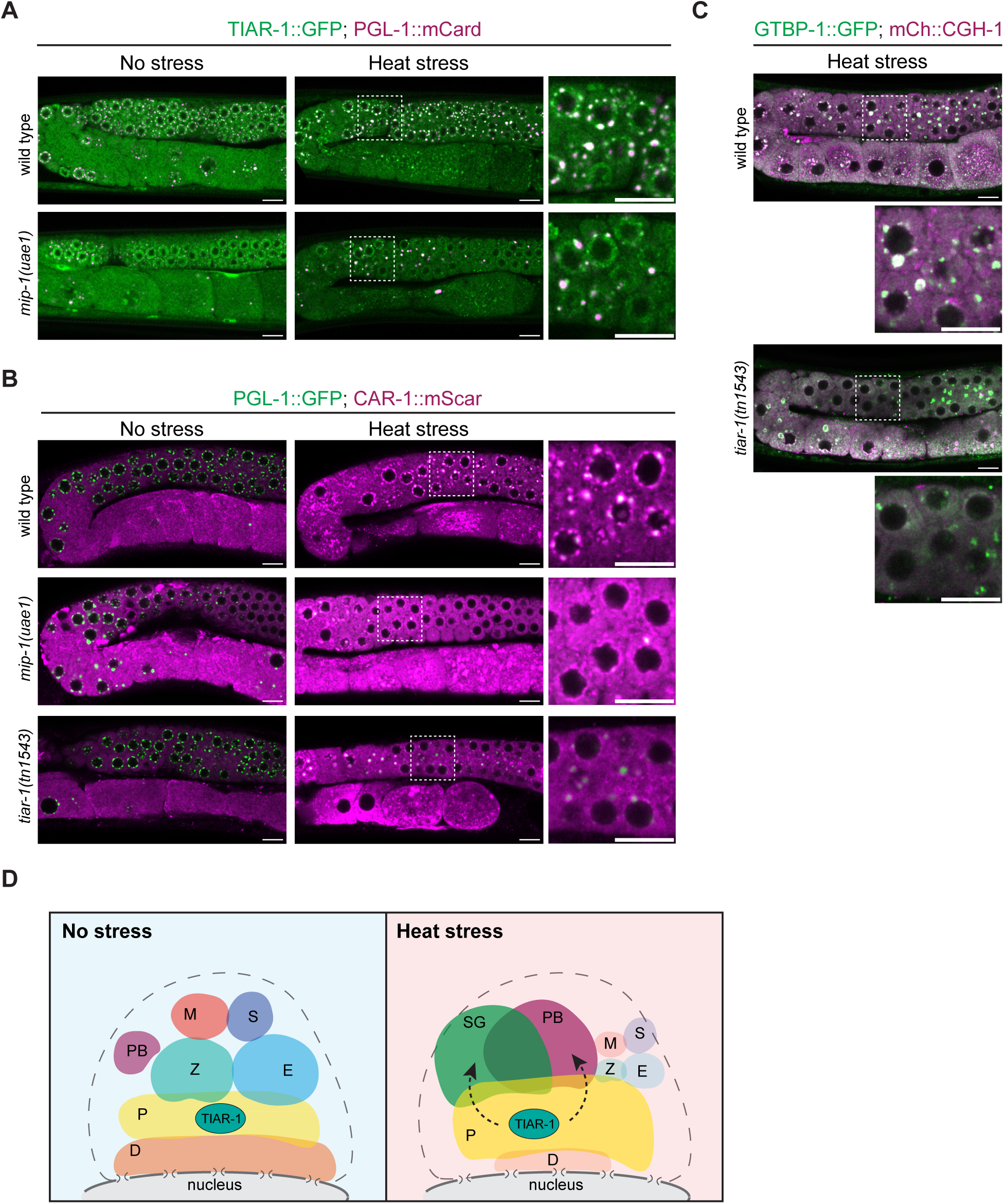
P granules serve as docking sites for stress granule and P-body assembly in germ cells under heat stress. (**A** - **C**) Representative live-fluorescent images of day one adult germlines co-expressing indicated markers colored by channel in wild type and indicated mutant backgrounds in no stress (20 °C) and/ or heat stress (31 °C, 3h). White dashed boxes outline the expanded image shown at the right (A-B) or below (C). Scale bars = 10µm. (**D**) Model illustrating the germ granule sub-compartment organization in the young adult germline in no stress (20 °C) and heat stress (31 °C, 3h). Abbreviations in cartoon P, Z, D, and E correspond to granule sub-compartments, (M) mutator focus, (S) SIMR focus, (PB) P-body, and (SG) stress granule. P granules are enlarged and promote SG and PB formation during heat stress. Stress granule factor TIAR-1 positioned within P granules facilitate the robust SG and PB assembly.

Since we found stress response factor TIAR-1 localizes to P granules, dependent on MIP-1, under normal and stress conditions, we wondered whether TIAR-1 may function at the interface of stress granule and P-body formation during heat stress. Like *mip-1(uae1)* mutants, we observed reduced perinuclear P granule and CAR-1 P-body formation in *tiar-1(tn1543)* mutant germlines compared to wild type (Fig. 8B). Further, under heat stress, we found a reduction in stress granules as marked by core stress granule assembly factor GTBP-1 concurrent with decreased CGH-1 P-body formation in *tiar-1(tn1543)* mutant germlines compared to wild type (Fig. 8C). This suggests that following heat stress, TIAR-1 may function at the P granule to promote stress granule and P-body formation (Fig. 8D). Together, our results provide insight into the distinct assembly of stress-responsive condensates in germ cells, where P granules contain stress response factors, like TIAR-1, enabling a robust and rapid response to stress (Fig. 8D). Overall, we uncover an important and highly adaptive role for germ granules in germ cell stress resilience.

## Discussion

Animals experiencing environmental stress must coordinate a stress response to protect normal cellular functions. In this study, we discovered a critical role for germ granules during heat stress in promoting stress granule and P-body assembly to protect germ cell viability. While we found germ granules undergo rapid remodeling under heat stress, including disrupted enrichment of most sub-compartment factors, P granule compartment factors, including PGL-1, remained highly intact. Under heat stress, stress granules and P-bodies formed at the cytoplasmic face of P granules, largely dependent on P granule assembly factors GLH-1/VASA helicase and LOTUS-domain protein MIP-1. Moreover, we found P granule assembly factors, GLH-1 and MIP-1, like conserved stress granule factors GTBP-1/G3BP1 and TIAR-1/TIA1, were critical for the recovery of germ cell viability after heat stress. We propose that germ granules function as a central stress-responsive hub to integrate the stress response in germ cells.

Intriguingly, while P granule compartment factors, such as PGL-1 and WAGO-1, largely maintained germ granule enrichment, we found heat stress induced an adaptive remodeling of most other germ granule sub-compartments. This remodeling included the reduced enrichment of several factors, including MUT-16, PRG-1, and WAGO-4, essential for functional RNA interference, piRNA-mediated gene silencing, and epigenetic inheritance ^14,54,55^. Our lab previously identified an interaction between germ granules and P-bodies in adult germ cells under normal growth conditions. P-body factors CGH-1 and CAR-1 were found to promote the compartmentalization of germ granules necessary for the robust inheritance of small RNA-based gene silencing ^39^. This same work identified Z granule factor Argonaute WAGO-4 as the link to the transgenerational silencing defect observed in *cgh-1* mutants. Here we observed disrupted Z granule localization of WAGO-4, as well as a significant dispersal of the Mutator foci assembly factor MUT-16 after heat stress. Components of these granules perform well-described roles in RNAi-triggered gene silencing and inheritance ^14,54,55^. Our findings suggest the specific disruption of these granules may contribute to the reported RNAi and transgenerational silencing defects observed after heat stress ^28^. In line with our observations, animals exposed to environmental stress reportedly showed a drastic reduction in Mutator foci assembly factor MUT-16-dependent small RNAs compared to control animals ^28^. Together, our findings highlight that heat stress contributes to dramatic remodeling in germ granule organization that are likely linked to the reduced small RNA populations and gene regulatory function.

Stress granules and P-bodies are clearly distinguishable RNA granules known to store mRNAs in animals but also share many components. During stress, they often exist in proximity to each other, suggesting they may coordinate mRNA processing, storage, and degradation. As observed in earlier reports, we similarly found P-bodies and stress granules highly co-localized after heat stress in *C. elegans* germ cells ^34–37^. Interestingly, previous studies identified that the mammalian ortholog of CGH-1, DDX6, promoted the formation of both P-bodies and stress granules in cells exposed to stress ^50^. This finding revealed that in certain conditions, stress granule assembly depends on P-bodies. However, in our investigation within germ cells following heat stress, we found stress granule and P-body formation is largely independent, more consistent with models of P-body and stress granule docking ^47,56–58^. Specifically, stress granule assembly was intact and perhaps even more robust in germ cells during knockdowns of core P-body assembly factors CGH-1 or CAR-1, while P-body assembly continued in the absence or knockdown of stress granule factors GTBP-1 and TIAR-1. Instead, our findings reveal a distinct organization principle in the *C. elegans* germline, where a third RNA condensate—the germ granule—acts as a scaffold that promotes the assembly of both stress granules and P-bodies during heat stress.

Germ granules perform well-established roles in germ cell development and inheritance ^1–5^. Here, we uncover a crucial role for germ granules in protecting germ cells from acute heat stress. First, our study revealed that P-bodies, and stress granules assemble at the cytoplasmic site of P granules after exposure to heat stress. This position suggested P granules may serve as seeding sites for stress granule and P-body assembly during the stress response. Indeed, loss of key germ granule assembly factors GLH-1 and MIP-1 lead to highly compromised stress granule and P-body assembly. This finding suggests the docking of the P-body and stress granule factors at the nuclear periphery may require interactions with either germ granule enriched RNAs or specific protein components. While stress granules, P-bodies, and germ granules have previously been observed in proximity together within the germline ^27,34–37^, whether their interaction performed a protective role during the stress response remained unclear. Here we observed that P granule assembly mutants exhibit defects in germ cell recovery from heat stress, closely resembling the defects observed in stress granule assembly mutants. Intriguingly, we showed that conserved stress granule factor TIAR-1 localizes to P granules in normal and stressful environments, suggesting it may be involved in mediating P granule and stress granule interactions ^35,42^. We propose that germ granules function as a central stress-responsive hub that enables germ cells to dynamically reorganize their RNA regulatory landscape during stress, coordinating the assembly and function of stress granules and P-bodies. Such remodeling may reflect a broader adaptation of the germline in which the allocation of RNA regulatory resources is reprioritized during acute stress—from processes that normally support small RNA-mediated silencing and inheritance toward mechanisms that promote stress adaptation and preserve germline function. By coordinating these RNA regulatory compartments, germ granules may therefore help germ cells balance their essential role in transmitting genetic information with the immediate need to survive and recover from environmental stress.

## Methods

### Caenorhabditis elegans strains and maintenance

Animals were grown on standard nematode growth media (NGM) plates seeded with the Escherichia coli OP50 strain at 20 °C. The strains used in this study are listed in Supplemental Material, Table S2.

### Heat stress assay

Day before heat stress 50-100 L4 stage worms were transferred to new plates. Next day 20-30 worms per condition were transferred to a 2% agarose pad on slide with 40 mM sodium azide. For no stress condition, animals kept on slide at room temperature for 15 minutes before imaging. For 37 °C 15 minute heat stress group, slides were placed on aluminum block kept inside 37 °C incubator. Slides were imaged immediately after 15 minutes on a Zeiss compound microscope or Zeiss LSM800 confocal microscope with a Plan-Apochromat ×40/1.4 Oil DIC M27 objective.

For 31 °C 3-hour heat stress condition, the day before heat stress 50 L4 stage worms were transferred to new plates in duplicate. Next day, one plate was wrapped with parafilm to make airtight. Plates were submerged in temperature-controlled water bath held at 31 °C for 3 hours. The no stress animals were kept at normal 20 °C growth temperature. For both no stress and heat stress conditions, 25 worms were transferred to a 2% agarose pad on slide with 40 mM sodium azide and immediately imaged on a Zeiss LSM800 confocal microscope with a Plan-Apochromat ×40/1.4 Oil DIC M27 objective

### RNA interference

RNAi was performed by feeding animals with *E. coli* HT115 (DE3) strains expressing the appropriate dsRNA. RNAi bacterial strains were obtained from the Ahringer *C. elegans* RNAi Collection (Source BioScience). Bacterial cultures were grown in Luria broth supplemented with 100 μg/mL ampicillin and 50 μg/mL tetracycline overnight at 37°C. Cultures were seeded on NGM plates containing 100 μg/mL ampicillin, 50 μg/mL tetracycline, and 1 mM IPTG and incubated at room temperature for 24 hours. HT115 (DE3) expressing empty RNAi vector L4440 was used for control. RNAi conditions were seeded in duplicate and fifty L1 stage animals were transferred to each plate. After three days, young adult animals were used in the heat stress assay described above.

### Confocal live imaging

Tagged fluorescent proteins were visualized in living nematodes by mounting young adult animals on 2% agarose pads with 40 mM sodium azide. Fluorescent images used for localization studies were captured using a Zeiss LSM800 confocal microscope with a Plan-Apochromat x40/1.4 Oil objective.

### Image processing and analysis

Image processing and quantification of fluorescent puncta were performed using ImageJ. Single slice images of gonads were used for quantification. ROIs were selected, and areas of ROIs were measured. For quantitative co-localization analysis, all image manipulations were performed with ImageJ using the Coloc 2 plugin. Images of 7–10 gonads were collected and quantified. Data were compared by two-tailed p values were calculated using Mann-Whitney-Wilcoxon test.

### Heat stress embryonic viability assay

Day before heat stress assay fifty L4 stage worms were transferred to new plate. On the next day, twenty-five worms per genotype were transferred to new plate. Animals remaining on the original plate were used as control (no stress) animals. For heat stress condition, these new plates were wrapped with parafilm to make airtight. Plates were submerged in temperature-controlled water bath held at 31 °C. After 3 hours, ten worms were transferred to individual plates at 20 °C normal growth temperature. For control (no stress) animals, ten worms from the original plate kept at 20 °C were transferred to individual plates at 20 °C normal growth temperature. After 8 hours, adult heat stress and control worms were transferred to new individual plates. Embryos laid during 0 to 8 hours after heat stress or controls are group 1. Total embryos laid per plate was recorded for group 1. After 12 more hours, adult heat stress and control worms were transferred to new single plate. Embryos laid during 8 to 20 hours after heat stress or controls are group 2. Total embryos laid per plate was recorded for group 2. After 24 more hours, heat stress or control worms were transferred to new single plate. Embryos laid during 20 to 44 hours after heat stress or controls are group 3. Total embryos laid per plate was recorded for group 3. For each group of embryos, 24 hours after parent worms were transferred off plates, hatched larvae were counted. Total number of lethal embryos were measured by subtracting number of hatched larvae from total embryos counted the day before. Unhatched embryos over total embryos laid are shown as percentage and labeled percentage of embryonic lethality.Complete experiment was performed twice.

### Statistics and Reproducibility

Data is expressed as mean values with error bars indicating the standard error mean values in all bar graphs or plots. Statistical significance was evaluated using two-tailed Student’s t-test or Mann-Whitney-Wilcoxon test where indicated. GraphPad Prism 10 or R scripts were used for statistical analysis. No statistical method was used to predetermine sample size. No data were excluded from the analyses. The experiments were not randomized. The Investigators were not blinded to allocation during experiments and outcome assessment. All data for this study were obtained from cultures of *C. elegans* animals at the specified larval or adult stages. All imaging experiments were conducted using the indicated number (n) of independent animals. All brood size and RNAi treatments were conducted using the indicated number (n) of independent animals.

## Acknowledgments

Some strains used in this study were provided by the Caenorhabditis Genetics Center (CGC) funded by NIH Office of Research Infrastructure Programs (P40 OD010440).

## Author contributions

Conceptualization: O.G. and H.-C.L., Investigation and data analyses: O.G., A.B., M.A., Writing: O.G. and H.-C.L.

