## Supplementary Figures for "Germ granules function as stress-responsive hubs to protect germ cell resilience"

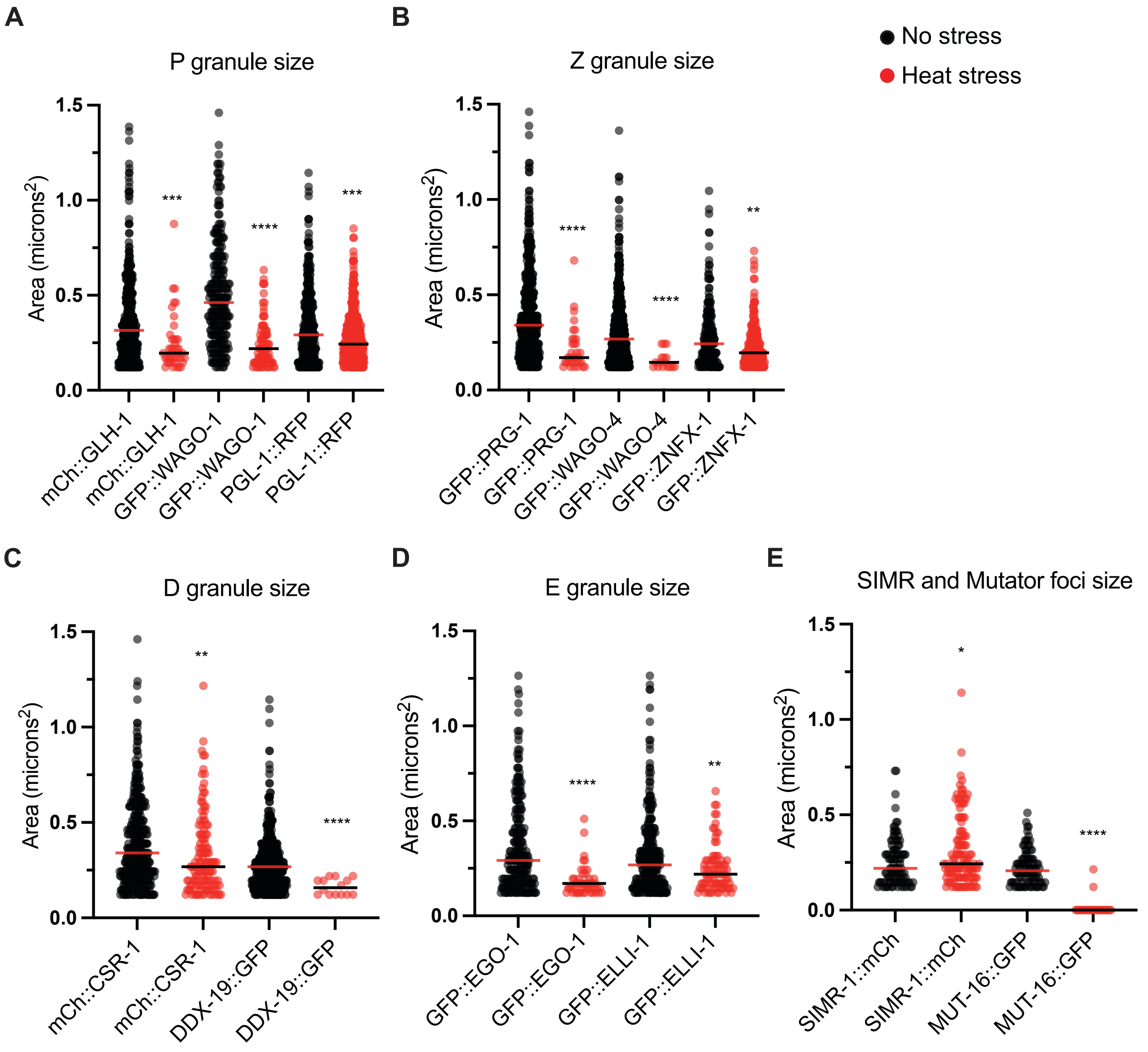


**Supplemental Figure 1: Germ granule compartments are mostly reduced in size after heat stress.**

**(A - E)** Plots depicting the granule size in area (microns$^2$) quantified for the indicated markers for N=8-12 independent germlines for no stress (20 °C) (black points) and heat stress (37 °C, 15m) (red points) conditions. Red and black lines mark the median value. Two-tailed p values were calculated using Mann-Whitney-Wilcoxon test. *, $p<0.05$, **, $p<0.01$; ***, $p<0.001$; ****, $p<0.0001$.

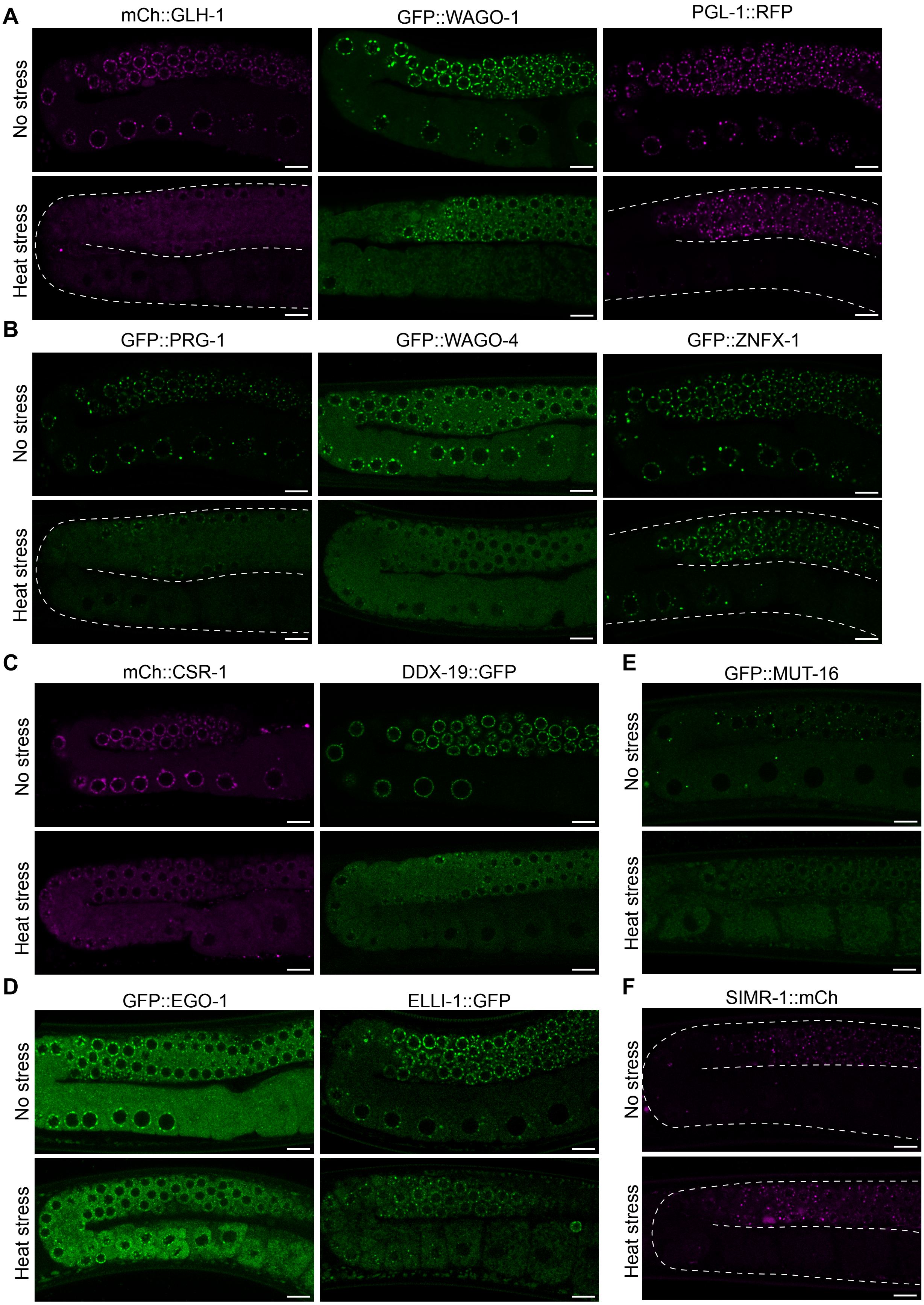
A
mCh::GLH-1
GFP::WAGO-1
PGL-1::RFP
No stress
Heat stress
B
GFP::PRG-1
GFP::WAGO-4
GFP::ZNFX-1
No stress
Heat stress
C
mCh::CSR-1
DDX-19::GFP
No stress
Heat stress
D
GFP::EGO-1
ELLI-1::GFP
No stress
Heat stress
E
GFP::MUT-16
No stress
Heat stress
F
SIMR-1::mCh
No stress
Heat stress

**Supplemental Figure 2: Germ granules throughout the germline are remodeled after heat stress.**

(**A** - **F**) Representative live-fluorescent images of young adult germlines expressing indicated granule compartment or foci markers labeled. Germlines of animals grown in no stress (20 °C) or animals exposed to heat stress (37 °C, 15 min). Compartments shown are P (**A**), Z (**B**), D (**C**), E (**D**), mutator foci (**E**), and SIMR foci (**F**). Dashed white line outlines germline. Scale bars = 10 μm.

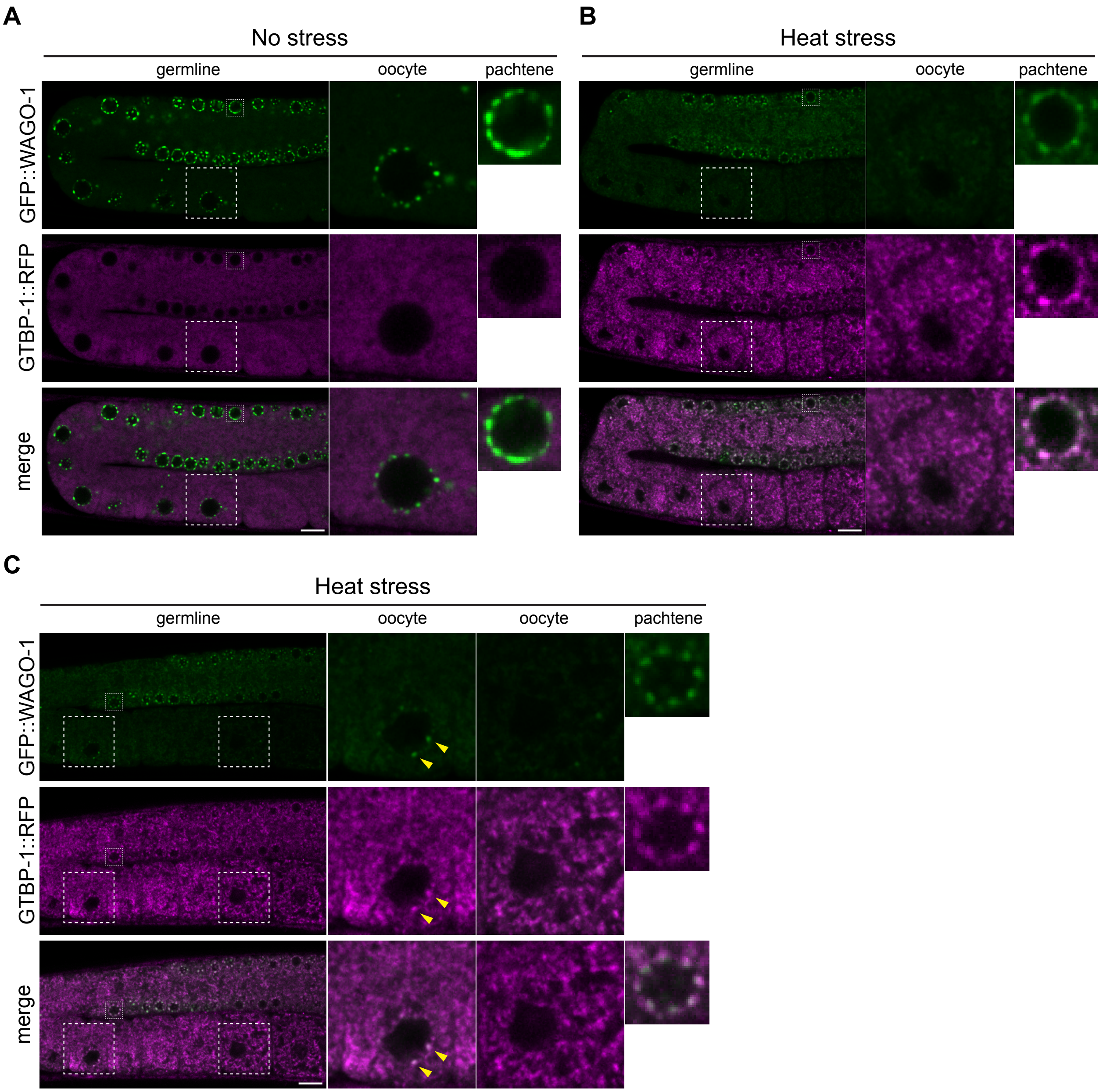


**Supplemental Figure 3: Stress granules form throughout the germline and with P granules after heat stress.**

(**A - C**) Representative live-fluorescent images of young adult germlines co-expressing GFP::WAGO-1 and GTBP-1::RFP for no stress (20 °C) (A) and two examples of heat stress (37 °C, 15m) (B and C). Large white dashed boxes outline oocyte nucleus enlarged. Small white dashed boxes outline pachytene nucleus enlarged. Yellow arrowheads point to co-localized perinuclear P granule and stress granule in oocyte. Scale bars = 10 μm.

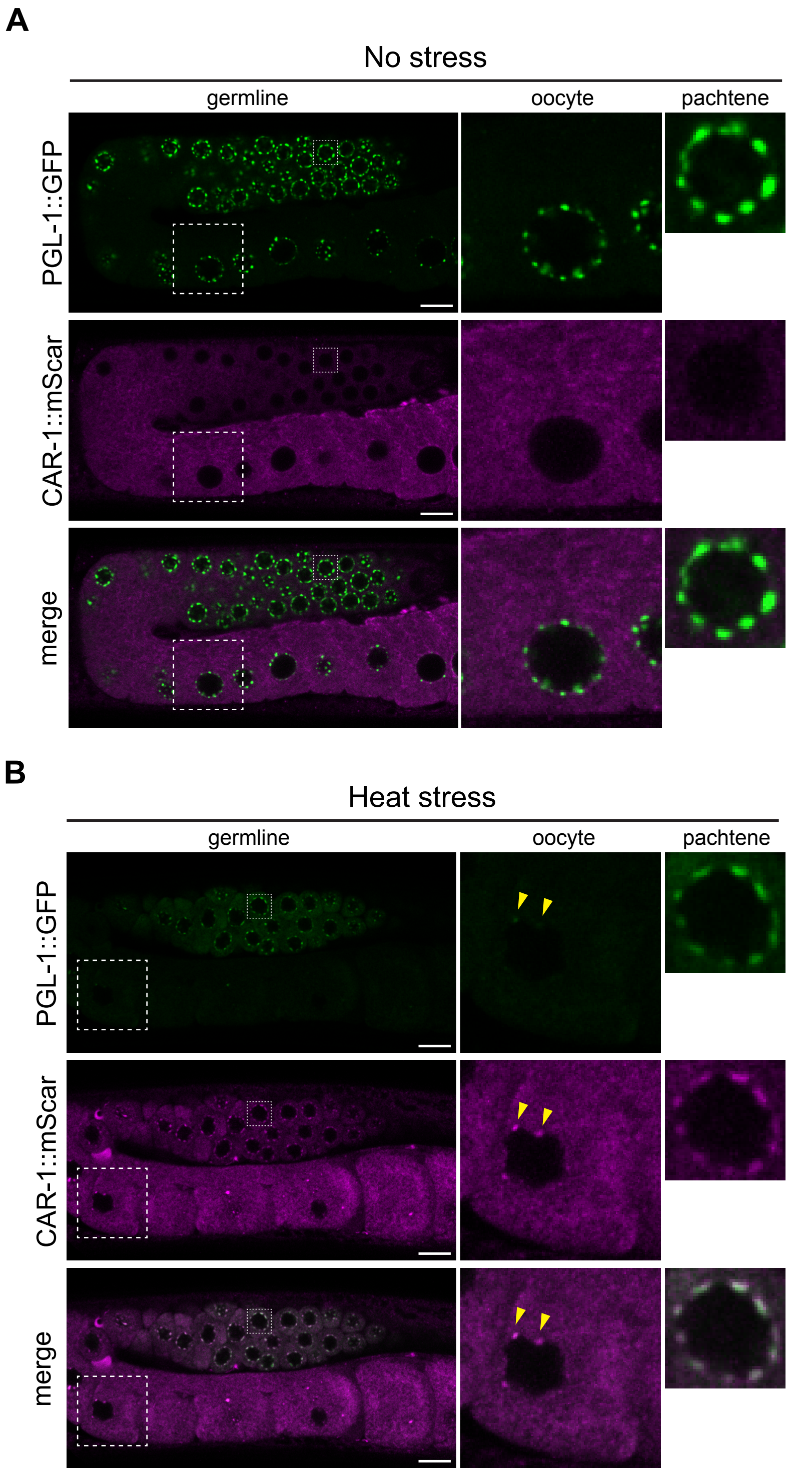


**Supplemental Figure 4: P-bodies form throughout the germline and with P granules after heat stress.**
(**A - C**) Representative live-fluorescent images of young adult germlines co-expressing PGL-1::GFP and CAR-1::mScarlet for no stress (20 °C) (A) and heat stress (37 °C, 15m) (B) conditions. Large white dashed boxes outline oocyte nucleus shown enlarged.

Small white dashed boxes outline pachytene nucleus shown enlarged. Yellow arrowheads point to co-localized perinuclear P granule and P-bodies in oocyte. Scale bars = 10 μm.

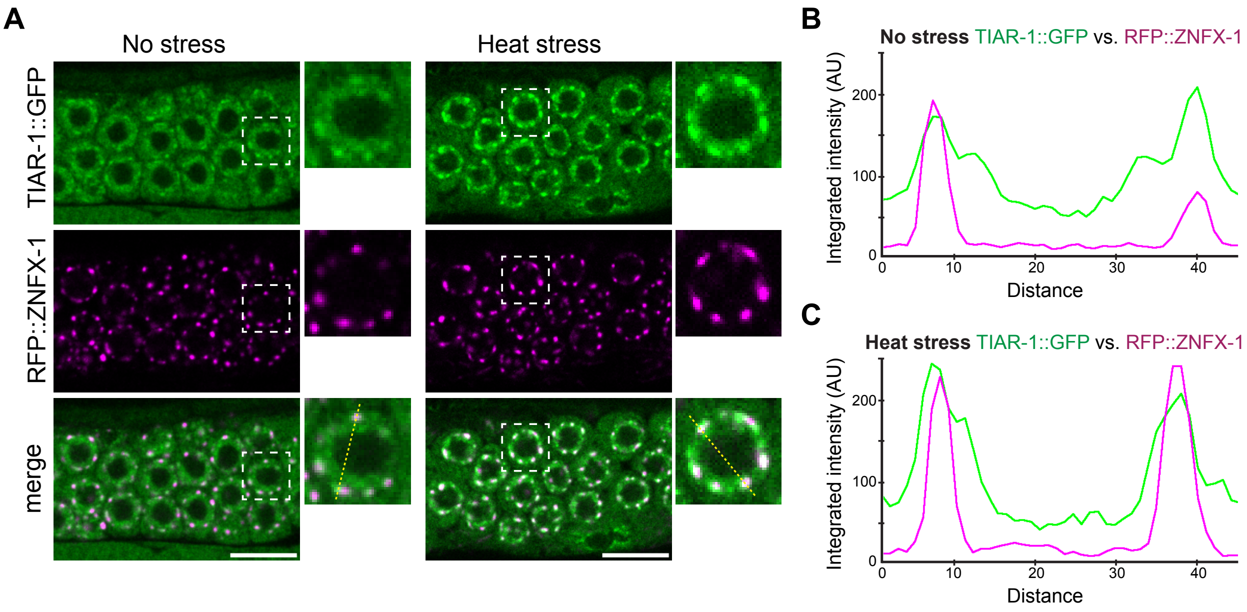


**Supplemental Figure 5: Stress granule factor TIAR-1 localizes adjacent to Z granules.**

(**A**) Representative live-fluorescent images of pachytene germ cells co-expressing TIAR-1::GFP and Z granule marker RFP::ZNFX-1 in no stress (20 °C) or heat stress (37 °C, 15 min) conditions. Dashed white boxes outline the individual germ cell nucleus shown in the enlarged image. Scale bars = 10 μm. (**B** and **C**) Line scans display the integrated fluorescent intensity corresponding to the yellow dashed line across the nucleus shown in (A) for the indicated markers labeled in green and magenta in no stress (B) and heat stress (C) conditions.

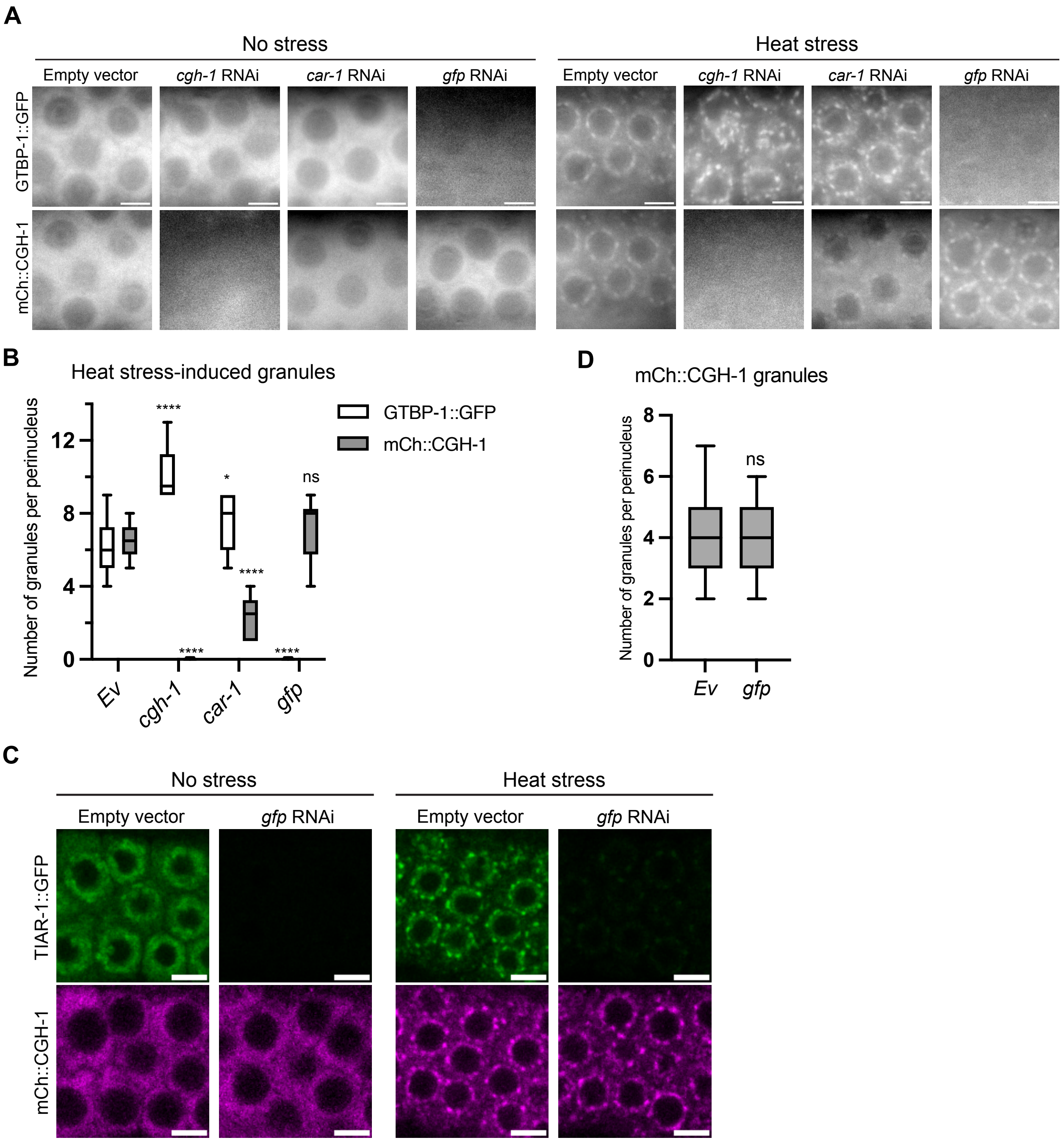


**Supplemental Figure 6: Perinuclear stress granules and P-bodies form largely independently in germ cells after heat stress.**

(**A**) Representative live-fluorescent images of young adult pachytene germ cells co-expressing stress granule marker GTBP-1::GFP and P-body marker mCh::CGH-1 for empty vector, *gfp*, *cgh-1*, or *car-1* RNAi conditions for no stress (20 °C) or exposure to heat stress (37 °C, 15m). Scale bars = 5 μm. (**B**) Boxplots showing the number of granules per germ cell perinucleus for indicated markers in animals exposed to heat stress (37 °C, 15m). Quantified for 5-6 cells from N=8-10 animals. (**C**) Representative live-fluorescent images of young adult pachytene germ cells co-expressing stress granule marker TIAR-1::GFP and P-body marker mCh::CGH-1 for empty vector or *gfp* RNAi conditions for no stress (20 °C) or exposure to heat stress (37 °C, 15m). Scale bars = 5 μm. (**D**) Boxplots showing the number of mCh::CGH-1 marked granules per germ cell perinucleus in animals exposed to heat stress (37 °C, 15m). Quantified for 5-6 cells from N=9 animals. For all boxplots, the line indicates the median value, the box indicates the first and third quartiles, and the whiskers indicate the min and max values. Two-tailed p values were calculated using Mann-Whitney-Wilcoxon test. *, $p<0.05$; ****, $p<0.0001$; ns, no significance.

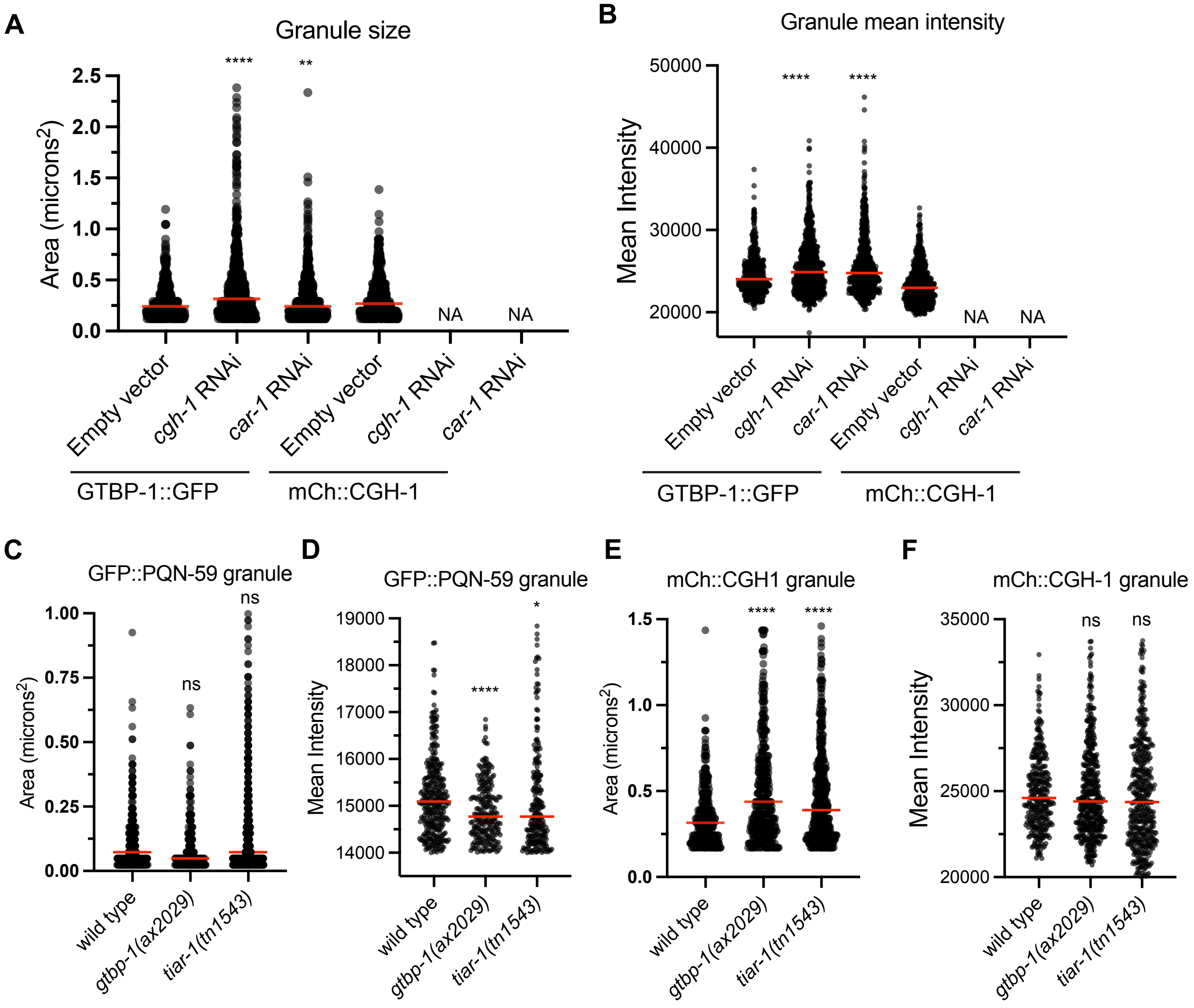


**Supplemental Figure 7: Perinuclear stress granule and P-body size and intensity after heat stress.**

(**A, C, E**) Plots depicting the threshold granule size in area (microns$^2$) quantified for the indicated markers in RNAi knockdown, or mutant conditions for heat stress (37 °C, 15m) animals. (**B, D, F**) Plots show the mean intensity per granule quantified for the indicated markers in RNAi knockdown, or mutant conditions for heat stress animals. (**A-F**) For all plots N=8-15 independent germlines. Red line marks the median value. NA indicates not available as no granules met the threshold. Two-tailed p values were calculated using Mann-Whitney-Wilcoxon test. **, p<0.01; ****, p<0.0001; ns, no significance.

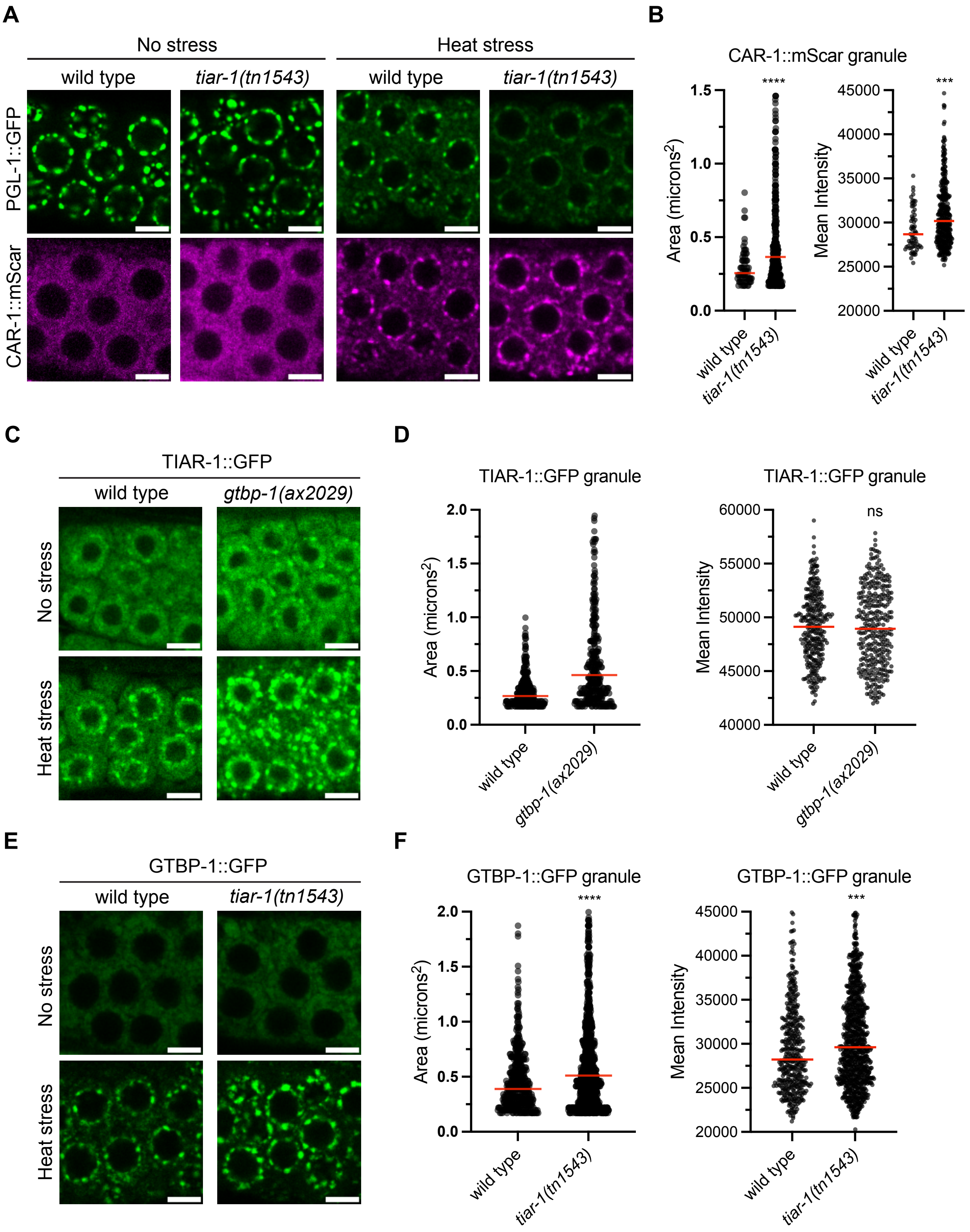


**Supplemental Figure 8: Role of stress granule factors in heat-induced granule assembly in germ cells.**

(**A, C, E**) Representative live-fluorescent images of young adult pachytene germ cells expressing the indicated markers in wild type and mutant backgrounds in no stress (20 °C) or exposure to heat stress (37 °C, 15m). Scale bars = 5 μm. (**B, D, F**) Plots depicting the threshold granule size in area (microns$^2$) quantified for the indicated markers in wild type or mutant conditions for heat stress (37 °C, 15m) animals (left). Plots show the mean intensity per granule quantified for the indicated markers in wild type or mutant conditions for heat stress (37 °C, 15m) animals (right). For all plots N=8-15 independent germlines. Red line marks the median value. Two-tailed p values were calculated using Mann-Whitney-Wilcoxon test. ***, $p<0.001$; ****, $p<0.0001$; ns, no significance.

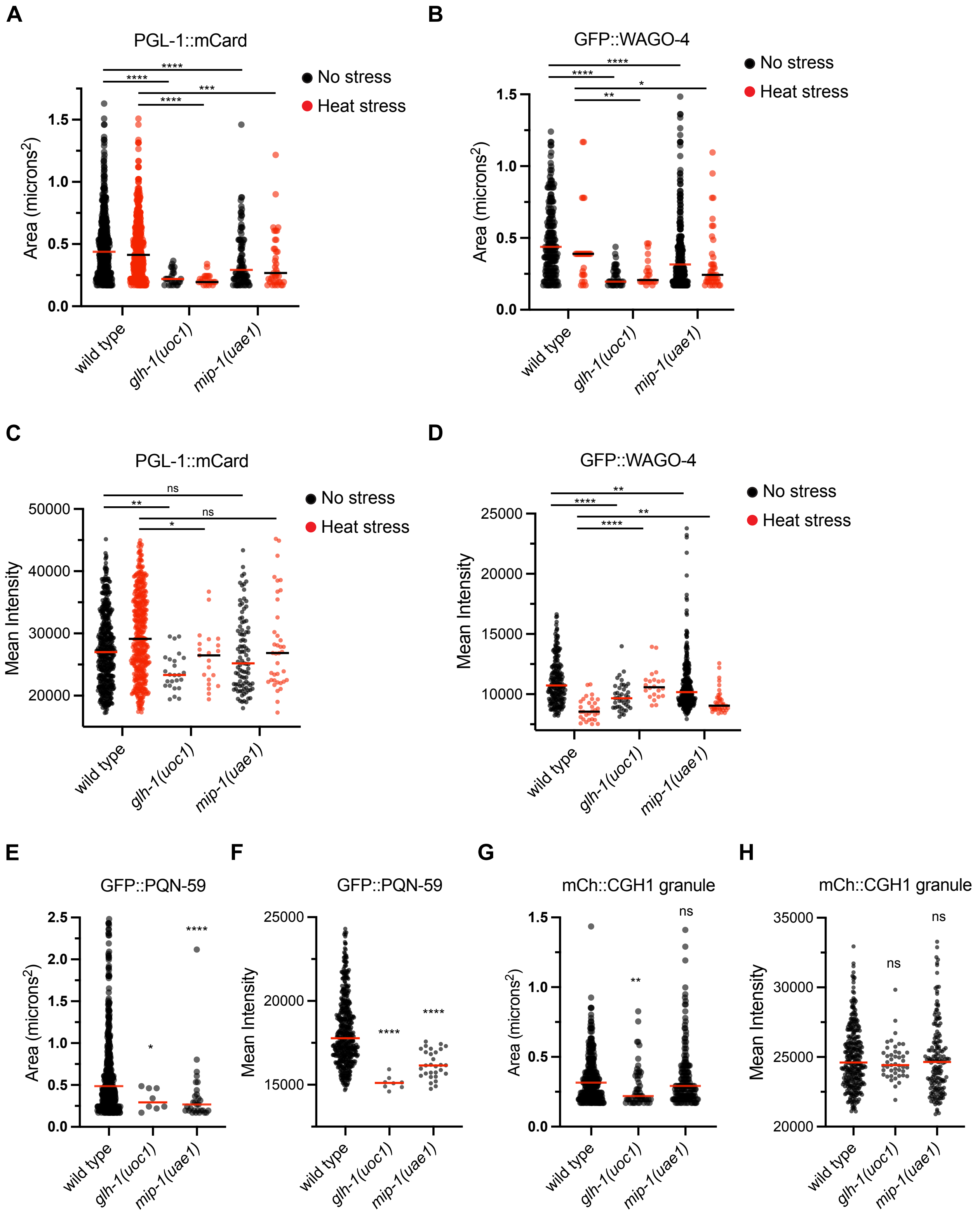


**Supplemental Figure 9: Germ granule assembly factors promote SG and P-body assembly at the germ cell perinucleus after heat stress.**

(**A** - **D**) Plots depicting the threshold granule size in area (microns$^2$) (A and B) and mean intensity (C and D) quantified for the indicated markers in wild type, *glh-1(uoc1)*, and *mip-1(uae1)* mutants in no stress (20 °C) (black points) or after heat stress (37 °C, 15m) (red points) for indicated markers. (**E** - **H**) Plots depicting the threshold granule size in area (microns$^2$) (E and G) and mean intensity (F and H) quantified for the indicated markers in wild type, *glh-1(uoc1)*, and *mip-1(uae1)* mutants after heat stress (37 °C, 15m) (black points) for indicated markers. All quantification for 5-6 cells from N=8-15 animals. Red and black lines mark the median value. Two-tailed p values were calculated using Mann-Whitney-Wilcoxon test. *, p<0.05; **, p<0.01; ***, p<0,001; ****, p<0.0001; ns, no significance.

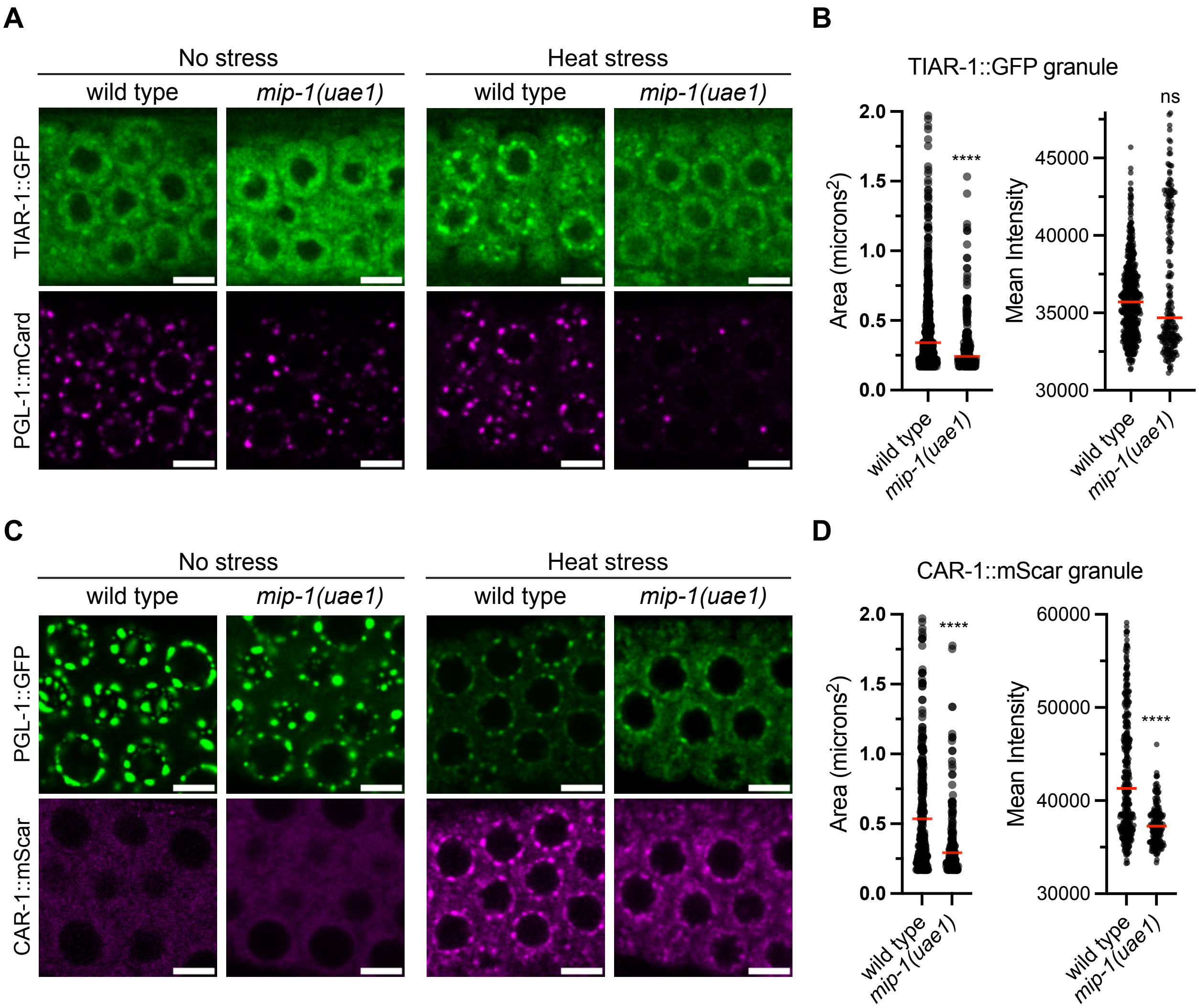


**Supplemental Figure 10: Germ granule assembly factor MIP-1 promotes SG and P-body assembly at the germ cell perinucleus after heat stress.**

(**A** and **C**) Representative live-fluorescent images of young adult pachytene germ cells expressing the indicated markers in wild type and mutant backgrounds in no stress (20 °C) or exposure to heat stress (37 °C, 15m). Scale bars = 5 μm. (**B** and **D**) Plots depicting the threshold granule size in area (microns$^2$) quantified for the indicated markers in wild type or mutant conditions for heat stress (37 °C, 15m) animals (left). Plots show the mean intensity per granule quantified for the indicated markers in wild type or mutant conditions for heat stress (37 °C, 15m) animals (right). For all plots N=8-15 independent germlines. Red line marks the median value. Two-tailed p values were calculated using Mann-Whitney-Wilcoxon test. ****, p<0.0001; ns, no significance.

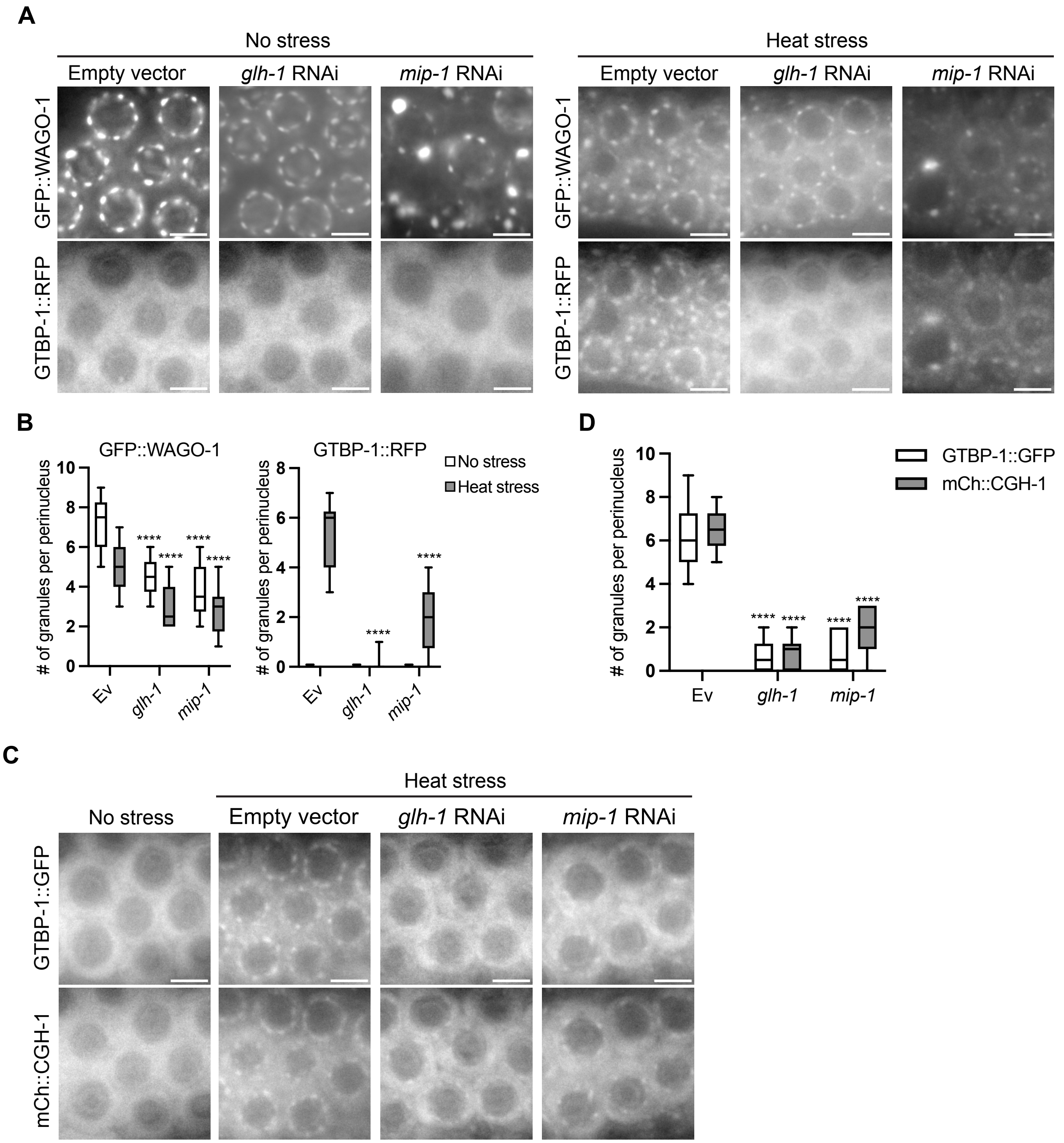


**Supplemental Figure 11: Germ granules promote heat stress-induced stress granule and P-body perinuclear formation.**

(**A**) Representative live-fluorescent images of young adult pachytene germ cells expressing both GFP::WAGO-1 and GTBP-1::RFP for control empty vector, *glh-1*, or *mip-1* RNAi conditions in no stress (20 °C) or heat stress (37 °C, 15m). Scale bars = 5 µm. (**B**) Boxplots showing quantification of the number of granules per germ cell perinucleus for experiment shown in (A). Ev, empty vector. No stress (white boxes), Heat stress (gray boxes). Quantified for 5-6 cells from N=6-9 animals. (**C**) Representative live-fluorescent images of young adult pachytene germ cells co-expressing GTBP-1::GFP and mCh::CGH-1 for empty vector, *glh-1*, or *mip-1* RNAi knockdown for heat stress (37 °C, 15m) conditions. All RNAi conditions were similar in no stress (20 °C). (**D**) Boxplots showing the number of granules per germ cell perinucleus in heat stress quantified from experiment (C) for the indicated markers. Ev, empty vector. For all boxplots, the line indicates the median value, the box indicates the first and third quartiles, and the whiskers indicate the min and max values. Two-tailed p values were calculated using Mann-Whitney-Wilcoxon test. ****, $p<0.0001$.

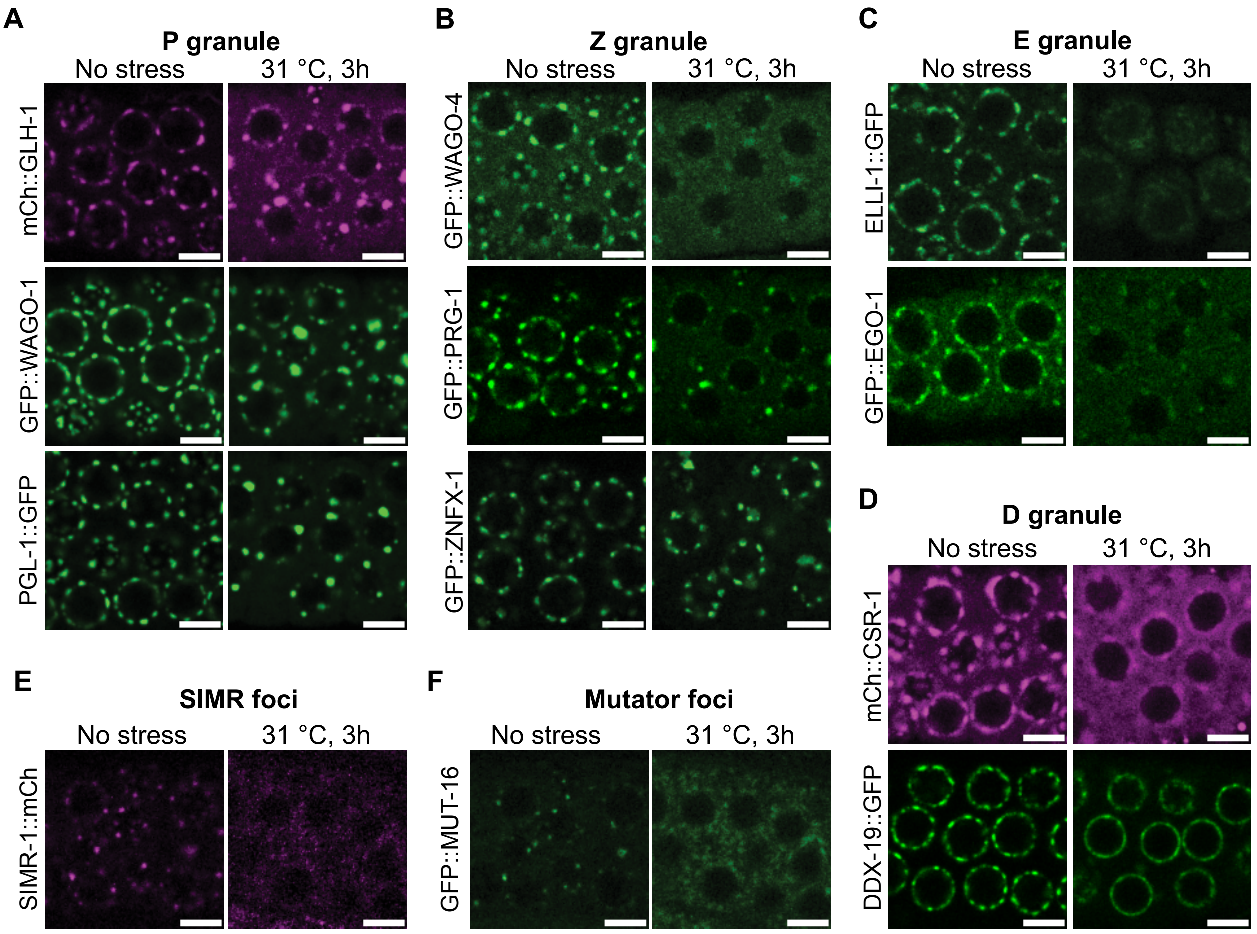


**Supplemental Figure 12: Germ granule compartments are remodeled after longer heat stress.**

(**A-F**) Representative live-fluorescent images of young adult pachytene germ cells expressing indicated granule compartment or foci markers labeled. Germlines of animals grown at 20 °C (No stress) or animals exposed to heat stress 31 °C, 3h. Scale bars = 5 µm.
