## Supplementary material for "Germ granules function as stress-responsive hubs to protect germ cell resilience": Table S1 and Table S2

**Table S1: Summary of granule compartment remodeling after heat stress.**

| Granule type | Marker | Normal growth (20 °C) | Heat stress (37 °C, 15m) | Heat stress (31 °C, 3h) |
|---|---|---|---|---|
| **Stress granule** | GTBP-1 | - | +++ | ++++ |
| | PQN-59 | - | +++ | ++++ |
| | TIAR-1 | + | ++ | ++++ |
| **P-body** | CGH-1 | - | +++ | ++++ |
| | CAR-1 | - | +++ | ++++ |
| | IFE-3 | - | +++ | NA |
| **P granule** | GLH-1 | +++ | + | ++++ |
| | WAGO-1 | +++ | + | ++++ |
| | PGL-1 | +++ | ++ | ++++ |
| **Z granule** | PRG-1 | +++ | + | +++ |
| | WAGO-4 | +++ | - | - |
| | ZNFX-1 | +++ | ++ | +++ |
| **E granule** | EGO-1 | +++ | + | - |
| | ELLI-1 | +++ | + | - |
| **D granule** | CSR-1 | +++ | + | + |
| | DDX-19 | +++ | + | + |
| **Mutator foci** | MUT-16 | ++ | - | - |
| **SIMR foci** | SIMR-1 | ++ | +++ | - |

Granule markers are organized by RNA granule type and germ granule sub-compartments. Markers scored for their observed granule size and intensity in pachytene germ cells of young adult animals at normal growth temperature (20 °C), heat stress (37 °C, 15m), and heat stress (31 °C, 3h). (-) indicates either no visible granules or granules did not meet the intensity threshold for quantification. (+ to ++++) serves as an increasing scale of granule size and intensity. NA, marker was not examined for that condition.

**Table S2: Strain list.**

| Strain | Genotype | Gene(s) | Description | Origin |
|---|---|---|---|---|
| N2 | *C. elegans wild isolate (Bristol)* | | Reference C. elegans. Variety Bristol. | CGC |
| GKC1 | *mip-1(uae1) III* | *mip-1* | | CGC |
| HCL118 | *glh-1(uoc1) I* | *glh-1* | | W Chen et al. 2020 |
| JH3176 | *gtbp-1(ax2029) IV* | *gtbp-1* | | CGC |
| JH3308 | *gtbp-1(ax5000[gtbp-1::tagRFP]) IV* | *gtbp-1* | | CGC |
| JH3199 | *gtbp-1(ax2055[gtbp-1::GFP]) IV* | *gtbp-1* | | CGC |
| DG4233 | *pqn-59(tn1737[GFP::3xFlag::pqn-59]) I* | *pqn-59* | | CGC |
| DG3922 | *tiar-1(tn1543[loxP::Cbr-unc-119(+)::loxP]) II* | *tiar-1* | | CGC |
| DG3922 | *tiar-1(tn1545[tiar-1::s::tev::GFP]) II* | *tiar-1* | | CGC |
| DZ419 | *GFP::wago-4 II; mCherry::cgh-1 III* | *wago-4; cgh-1* | Endogenously tagged by CRISPR-Cas9 editing in previous study | Z Du et al. 2023 |
| JH4186 | *car-1(syb8217[car-1::wrmScarlet] I* | *car-1* | | CGC |
| SHG682 | *ife-3(ust639[ife-3::GFP::3xFlag]) V* | *ife-3* | | CGC |
| HCL390 | *wago-1(tor113[GFP::3xFLAG::wago-1]) I; gtbp-1(ax5000[gtbp-1::tagRFP]) IV* | *wago-1; gtbp-1* | JMC219 crossed to JH3308 | This study |
| HCL391 | *mCherry::cgh-1 III; gtbp-1(ax2055[gtbp-1::GFP]) IV* | *cgh-1; gtbp-1* | DZ419 crossed to JH3199 | This study |
| HCL392 | *tiar-1(tn1545[tiar-1::s::tev::GFP]) II; pgl-1(gg640[pgl-1::3xflag::mCardinal]) IV* | *tiar-1; pgl-1* | DG3922 crossed to HCL366 (outcrossed from YY1492) | This study |
| HCL125 | *gfp::prg-1 flag::mCherry::glh-1 I* | *prg-1; glh-1* | | W Chen et al. 2020 |
| JMC219 | *tor113[GFP::3xFLAG::wago-1] I* | *wago-1* | | CGC |
| YY968 | *znfx-1(gg544[3xflag::gfp::znfx-1]) II; pgl-1(gg547[pgl-1::3xflag::tagRFP]) IV* | *znfx-1; pgl-1* | | CGC |

| | | | | |
|---|---|---|---|---|
| HCL155 | *ego-1[3xflag::gfp::ego-1] I* | *ego-1* | | W Chen et al. 2022 |
| SHG1682 | *elli-1(ust354[elli-1::GFP::3xFlag]) IV* | *elli-1* | | CGC |
| HCL376 | *wago-1(tor113[GFP::3xFLAG::wago-1]) I; csr-1(gc029[degron::mCherry::3xflag::ha::csr-1]) IV* | *wago-1; csr-1* | JMC219 crossed to MHE69 received from Germano Cecere lab | This study |
| SHG2207 | *ddx-19(ust399[ddx-19::GFP::3xFlag]) II* | *ddx-19* | | CGC |
| USC744 | *simr-1(cmp15[simr-1::mCherry::2xHA loxP Cbr-unc-119 loxP]) mut-16(cmp3[mut-16::gfp::3xFLAG + loxP]) I; unc-119(ed3) III* | *simr-1; mut-16* | | Received from Carolyn Phillips lab |
| HCL451 | *pqn-59(tn1737[GFP::3xFlag::pqn-59]) I; gtbp-1(ax2029) IV* | *pqn-59; gtbp-1* | DG4233 crossed to JH3176 | This study |
| HCL452 | *GFP::wago-4 II; mCherry::cgh-1 III; gtbp-1(ax2029) IV* | *wago-4; cgh-1; gtbp-1* | DZ419 crossed to JH3176 | This study |
| HCL453 | *pqn-59(tn1737[GFP::3xFlag::pqn-59]) I; mip-1(uae1) III* | *pqn-59; mip-1* | DG4233 crossed to GKC1 | This study |
| HCL454 | *glh-1(uoc1) I; GFP::wago-4 II; mCherry::cgh-1 III* | *glh-1; wago-4; cgh-1* | DZ419 crossed to HCL118 | This study |
| HCL455 | *GFP::wago-4 II; mCherry::cgh-1 mip-1(uae1) III* | *wago-4; cgh-1; mip-1* | DZ419 crossed to GKC1 | This study |
| HCL456 | *pqn-59(tn1737[GFP::3xFlag::pqn-59] car-1(syb8217[car-1::wrmScarlet] I* | *pqn-59; car-1* | DG4233 crossed to JH4186 | This study |
| HCL457 | *pqn-59(tn1737[GFP::3xFlag::pqn-59]) I; pgl-1(gg640[pgl-1::3xflag::mCardinal]) IV* | *pqn-59; pgl-1* | HCL453 crossed to HCL336 (outcrossed from YY1492) | This study |
| HCL459 | *pqn-59(tn1737[GFP::3xFlag::pqn-59]) I; mip-1(uae1) III; pgl-1(gg640[pgl-1::3xflag::mCardinal]) IV* | *pqn-59; mip-1; pgl-1* | HCL457 crossed to GKC1 | This study |
| HCL460 | *pqn-59(tn1737[GFP::3xFlag::pqn-59]) glh-1(uoc1) I; pgl-1(gg640[pgl-1::3xflag::mCardinal]) IV* | *pqn-59; glh-1; pgl-1* | HCL457 crossed to HCL118 | This study |
| JH3269 | *pgl-1(ax3122[pgl-1::gfp]) IV* | *pgl-1* | | CGC |
| HCL461 | *car-1(syb8217[car-1::wrmScarlet] I; pgl-1(ax3122[pgl-1::gfp]) IV* | *car-1; pgl-1* | JH3269 crossed to JH4186 | This study |

| | | | | |
|---|---|---|---|---|
| HCL462 | *pqn-59(tn1737[GFP::3xFlag::pqn-59]) I; tiar-1(tn1543[loxP::Cbr-unc-119(+)::loxP]) II* | *pqn-59; tiar-1* | DG4233 crossed to DG3929 | This study |
| HCL463 | *tiar-1(tn1545[tiar-1::s::tev::GFP]) II; gtbp-1(ax2029) IV* | *tiar-1; gtbp-1* | DG3922 crossed to JH3176 | This study |
| HCL464 | *car-1(syb8217[car-1::wrmScarlet] I; mip-1(uae1) III; pgl-1(ax3122[pgl-1::gfp]) IV* | *car-1; pgl-1; mip-1* | HCL461 cross to GKC1 | This study |
| HCL465 | *car-1(syb8217[car-1::wrmScarlet] I; tiar-1(tn1543[loxP::Cbr-unc-119(+)::loxP]) II; pgl-1(ax3122[pgl-1::gfp]) IV* | *car-1; pgl-1; tiar-1* | HCL461 cross to DG3922 | This study |
| HCL466 | *tiar-1(tn1545[tiar-1::s::tev::GFP]) II; gtbp-1(ax2029) IV* | *tiar-1; gtbp-1* | JH3308 crossed to DG3922 | This study |
| HCL467 | *tiar-1(tn1545[tiar-1::s::tev::GFP]) II; mip-1(uae1) III pgl-1(gg640[pgl-1::3xflag::mCardinal]) IV* | *tiar-1; mip-1; pgl-1* | GKC1 crossed to HCL392 | This study |
| HCL468 | *tiar-1(tn1543[loxP::Cbr-unc-119(+)::loxP]) II; gtbp-1(ax2055[gtbp-1::GFP]) IV* | *tiar1; gtbp-1* | DG3929 crossed to JH3199 | This study |
| HCL469 | *tiar-1(tn1545[tiar-1::s::tev::GFP]) II; pgl-1(gg640[pgl-1::3xflag::mCardinal]) IV* | *tiar-1; pgl-1* | DG3922 crossed to HCL366 (outcrossed from YY1492) | This study |
| HCL470 | *tiar-1(tn1545[tiar-1::s::tev::GFP]) II; mCherry::cgh-1 III* | *tiar-1; cgh-1* | DG3922 crossed to DZ419 | This study |
| HCL471 | *tiar-1(tn1545[tiar-1::s::tev::GFP]) znfx-1(gg634[HA::tagRFP::znfx-1]) II.* | *tiar-1; znfx-1* | DG3922 crossed to YY1446 | This study |
| HCL472 | *tiar-1(tn1543[loxP::Cbr-unc-119(+)::loxP]) II; mCherry::cgh-1 III; gtbp-1(ax2055[gtbp-1::GFP]) IV* | *tiar-1; cgh-1; gtbp-1* | DG3929 xrossed to HCL391 | This study |
